# Turning microglia neuroprotective: Towards connexin43-specific therapy of Alzheimer’s disease

**DOI:** 10.1101/2024.08.06.606883

**Authors:** Yixun Su, Hui Li, Wenjie Zhang, Shi Tao, Qi Wang, Mi Zhou, Yong Tang, Hui Chen, Alexei Verkhratsky, Zhengbao Zha, Jianqin Niu, Chenju Yi

## Abstract

Alzheimer’s disease (AD) is the major cause of senile dementia without effective therapeutic strategies. The fundamental role of microglia in AD pathology, particularly in the early stages, is well acknowledged, although cell-specific therapeutic targets were not identified. Here we show that microglial connexin 43 (Cx43) hemichannels controls microglial reactivity in AD, thus being a promising therapeutic target. We discovered a marked increase in Cx43 protein in the periplaque microglia in the post-mortem tissue from AD patients. Subsequently, using the APP_swe_/PS1_dE9_ mouse model of AD, we demonstrated that microglial Cx43 operating as hemichannels influences microglial function, which in turn affects β-amyloid pathology. Ablation of microglial Cx43 hemichannels by genetic knockout shifted microglia to neuroprotective phenotype, which promoted the microglia-plaque interaction while suppressing the neurotoxic microglial signature, thereby mitigating the progression of AD. Following this lead, we developed a novel formulation of a small molecule peptide, lipid nanoparticle-delivered molecule TAT-Cx43_266-283_ (TAT-CX43@LNPs), which selectively blocks Cx43 hemichannels. Our preclinical trial demonstrated its efficacy in delaying and rescuing β-amyloid-related neuropathology and cognitive impairment in AD mice. This study provides strong evidence to progress our novel drug into clinical trials and translate it to disease-preventing (when administered in the early disease stages) and disease-modifying agents.

## Introduction

Alzheimer’s disease (AD) is the most prevalent neurodegenerative and cognitive disorder among the elderly, with projections indicating doubling of AD-related dementia cases in Europe and tripling globally by 2050 ^1^. Decades of extensive research on AD were unable to fully reveal its complex and enigmatic pathophysiology ^2^. Ongoing new drug development efforts have been mainly guided by the widely accepted amyloid cascade hypothesis, which led to a staggering 98% failure rate in clinical trials ^3,4^. The recent β-amyloid (Aβ) antibody therapies are only suitable for AD patients at the very early stages, yet still failing to halt its progression ^5^. To address the stalemate in treating this irreversible disease, we must investigate new targets to tackle mechanisms beyond Aβ accumulation ^1^.

Among an array of cell types involved in early-stage AD ^6^, our attention is on microglia which are the first responders to the central nervous system (CNS) pathology, including AD ^7^. Microglial cells undergo significant and complex pathological changes at the pre-plaque stage in AD mice ^8^. These changes are also observed in patients with mild cognitive impairment that manifest the initial stages of AD ^9–11^. Such microglia was called disease-associated microglia ^12^, characterized by overexpression of genes (such as *Trem2*, *Apoe*, and *Grn*) associated with AD ^1,13,14^. Recent studies highlighted the dual role of microglia in AD. Microglial cells establish close interactions with Aβ plaques, forming a core-shell structure ^15^ to restrict the damage to the neighboring neuropil and actively participate in Aβ clearance ^13^. However, reactive microglia also contribute to neurodegeneration by engaging in aberrant synapse removal and inducing a pro-inflammatory, neurotoxic microenvironment ^13,16,17^. Although extensively studied, targeting microglia remains an uncharted area due to their remarkable flexibility and multiple roles during disease progression. Nevertheless, any cell-specific approach to shift microglia towards a neuroprotective state may represent a new approach in AD treatment.

A unique feature of pathological microglia during AD progression is associated with connexin 43 (Cx43). An upregulation of Cx43 in periplaque microglia in AD mice was demonstrated by *in situ* single-cell transcriptome mapping ^15^. Elevated Cx43 proteins were also confirmed by proteomic analysis of microglia isolated from AD mice ^18^. Connexins form gap-junction channels that are fundamental for cell-to-cell communication, and hemichannels for autocrine or paracrine signaling. Connexin expression represents a distinctive feature of macroglia in the CNS, connecting astroglia and oligodendroglia into panglial syncytia ^19^. Recent studies suggested elevated Cx43 expression and altered channel functions as the critical elements in AD pathology ^20–26^, while Aβ exposure has been found to activate microglial Cx43 hemichannels *in vitro* ^27^. However, the detailed contribution of microglial Cx43 in AD pathology and the potential of microglial Cx43 as a therapeutic target remain largely unexplored.

Our study is the first to explore a cell-specific treatment for AD, by targeting Cx43 hemichannels. Our in-depth examination of post-mortem human AD brains confirmed an abnormal increase in microglial Cx43. Using the APP_swe_/PS1_dE9_ (APP/PS1) mouse model of AD, we further identified microglial Cx43-related pathology as a novel mechanism in AD pathogenesis, while demonstrating that ablation of microglial Cx43 hemichannels shifted microglia to a neuroprotective reactive state. We translated this finding into a novel formulation, a lipid nanoparticle-based drug delivery system, which inhibits microglial Cx43 hemichannels to arrest and reverse AD progression in the pre-clinical trial.

## Results

### Increased microglial Cx43 in AD patient brains

We first analyzed the global changes in Cx43 expression in AD patient brains from online transcriptome and proteome databases. Analysis of the ROSMAP-DLPFC bulk RNA-sequencing data ^28^ showed that Cx43 RNA expression is significantly increased in AD patients and positively correlates with the Braak stage (**Fig 1A, Fig S1A**). Similarly, data from a multi-center consortium proteomics study (ACT-BLSA-Banner-MSSB) ^29^ revealed a positive correlation between Cx43 protein level and the Braak stage, indicating a significant elevation of Cx43 in AD patients compared to the healthy controls and asymptomatic AD patients (defined as cognitively normal individuals with AD neuropathological hallmarks ^30^) (**Fig 1B**). However, the cell-specific distribution of elevated Cx43 remained unknown.

**Fig 1.**
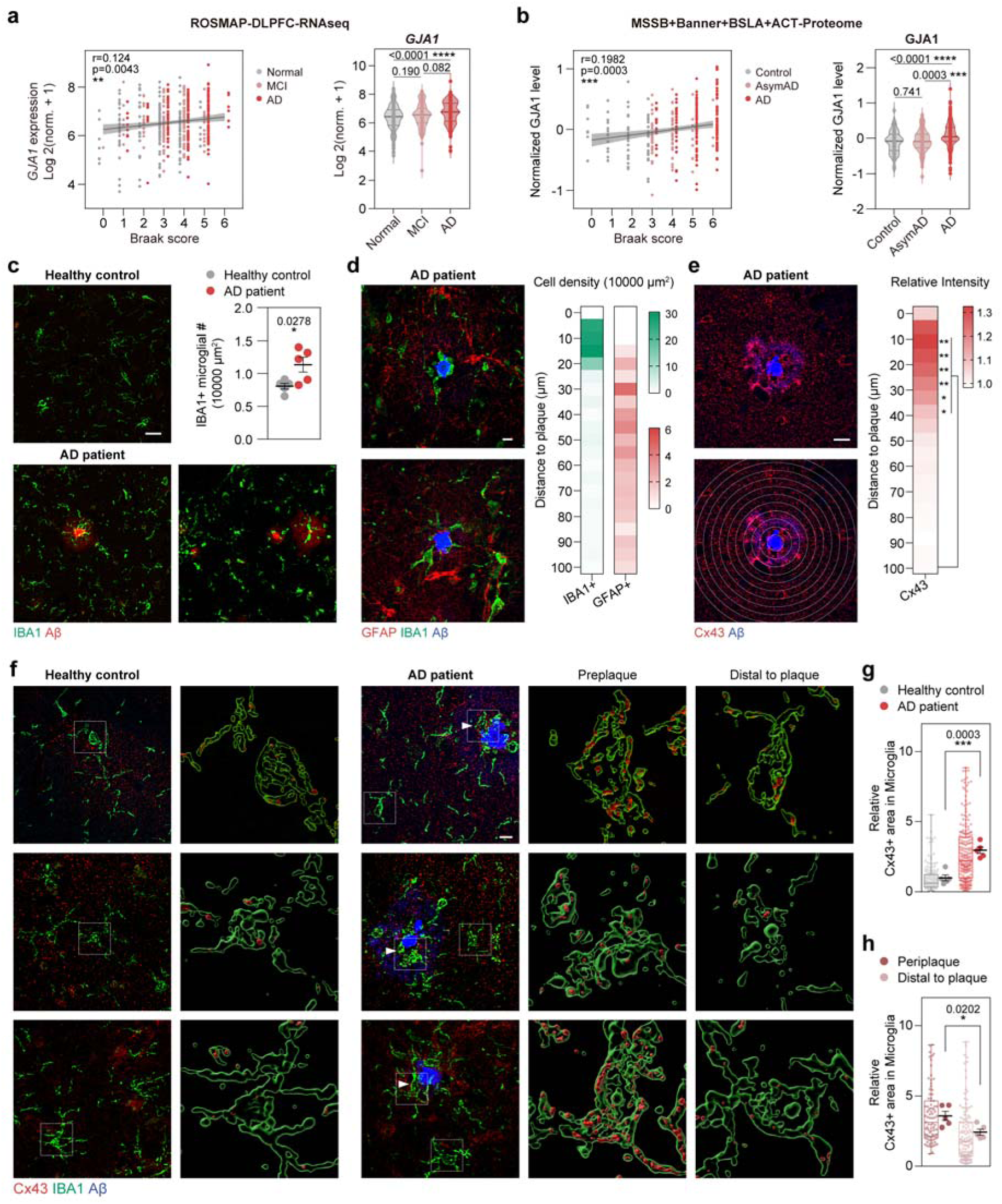
Increased microglial Cx43 in the brains from AD patients. (A) Cx43 (gene name: *GJA1*) expression from bulk RNA-sequencing of the human dorsolateral prefrontal cortex in the ROSMAP cohort was plotted against the Braak score, revealing a positive correlation. Right: expression of Cx43 in healthy, mild cognitive impairment (MCI), and AD group. (B) Cx43 protein level from a proteomic study of a multicenter cohort (Baltimore Longitudinal Study of Aging (BLSA), Banner Sun Health Research Institute (Banner), Mount Sinai School of Medicine Brain Bank (MSSB), Adult Changes in Thought Study (ACT)) was plotted against the Braak score. Right: Cx43 protein level in control, asymptomatic AD (asymAD), and AD group. (C) Representative imaging of IBA1 and Aβ immunostaining of healthy controls and AD patients, scale bar: 10 μm. Right: Quantification of IBA1+ microglia number. N = 5 subjects. (D) Representative images of IBA1, GFAP, and Aβ immunostaining of AD patients, scale bar: 10 μm. Right: Distribution of microglia and astrocytes at different distances from Aβ plaques. Cells around total of 30 plaques were calculated. (E) Immunostaining of Cx43 and Aβ, scale bar: 10 μm. Cx43 staining intensity at different distances from Aβ plaques (n=23) was quantified and compared to the area distal to plaque (55-100 μm). (F) Representative images of Cx43, IBA1, and Aβ immunostaining of healthy controls and AD patients, scale bar: 10 μm. The area within white boxes was 3D rendered by Imaris, displaying Cx43 signal in IBA1+ domain. Arrowhead: Periplaque microglia. (G) Quantification of Cx43+ area within IBA1+ domain in normal controls and AD patients, normalized to the mean of normal controls. N = 5 subjects, n = 145 (control) and 207 (AD) cells. (H) Quantification of Cx43+ area within IBA1+ domain in periplaque microglia and distal microglia in AD patients, normalized to the mean of normal controls. N = 5 subjects, n = 85 (periplaque) and 121 (distal) cells. Dot plots show mean ± SEM, each data point, and *p*-value. Significance: * *p* < 0.05, ** *p* < 0.01, *** *p* < 0.001, **** *p* < 0.0001.

Thus, we performed immunostaining on five post-mortem human AD brains and five age- and sex-matched healthy control brains **(Fig S1B)**. We found a significant increase in microglia numbers in the AD brain, microglial density was the highest in the vicinity of Aβ plaques (**Fig 1C-D**). We further analyzed the Cx43 protein level in relation to Aβ plaques, and found that Cx43 expression was also significantly higher in the periplaque area (**Fig 1E**). To evaluate the changes in microglial Cx43 levels, we quantified the colocalization of Cx43 with the microglial marker IBA1 using super-resolution microscopy. We found that the Cx43+ area associated with microglia was increased by 194 ± 31% (N = 5 subjects, *p* = 0.0003) in the AD patient brains when compared to the healthy controls (**Fig 1F-G**). It was suggested that microglia at the adjacent areas of Aβ plaques constitute a subset of disease-associated microglia with distinctive morphological and transcriptional changes, which may play a significant role in Aβ plaque formation, containment, and neurodegeneration ^12,15^. We therefore classified microglia into two groups: the periplaque microglia with their cell body located within 50 μm off the plaque center and in contact with Aβ plaque, and the distal microglia located > 50 μm to the plaque center. We found that the level of Cx43 was 46.0 ± 15.9% (N = 5 subjects, *p* = 0.0202) greater in periplaque microglia when compared to the distal microglia (**Fig 1F, H**).

Pannexin 1 (Panx1) is a plasmalemmal channel distantly related to the connexin family; Panx1 is expressed in microglia and can be affected by neurodegeneration ^31^. We analyzed Panx1 in post-mortem brain samples and found that Panx1 expression was low in microglia, and was not significantly altered in AD brains (**Fig S1C**). As a reference, we also quantified Cx43 changes in the GFAP+ astrocytes in controls and AD patients. Astrocytic Cx43 level was also significantly increased in AD patient brains, but the degree of the change (49.1 ± 18.8% compared to the control, N = 5 subjects, *p* = 0.0310) was lower compared to that of microglia (**Fig S1D**).

Taken together, these observations suggest that a marked increase in Cx43 is a specific feature characterizing pathological microglia in AD.

### Increased microglial Cx43 and hemichannel activity correlates with A**β** pathology in a mouse model of AD

We used the APP/PS1 mouse model, known to resemble human AD-related amyloid pathology in distinct disease stages (2 months: pre-onset; 4 months: onset of Aβ pathology; 9 months: profound Aβ pathology and measurable cognitive decline) ^32,33^ **(Fig S2A)**. In APP/PS1 mice, microglial density was high in the periplaque area **(Fig S2B)**. We first accessed the immunofluorescence staining of Cx43 in the hippocampus, which is known to regulate cognitive function and is impaired by AD ^34,35^. There were no significant changes in Cx43 levels in the hippocampus of 2- and 4-month-old APP/PS1 mice. However, a marked increase in Cx43 immunoreactivity was evident in 9-month-old APP/PS1 mice (**Fig 2A**). Further analysis revealed an increase in the Cx43 immunoreactivity in the periplaque areas in both 4 and 9-month-old mice, and an even marked increase in Cx43 staining intensity was observed in the periplaque areas in 9-month-old mice (**Fig 2B**), suggesting the increase in Cx43 levels is closely associated with Aβ plaques.

**Fig 2.**
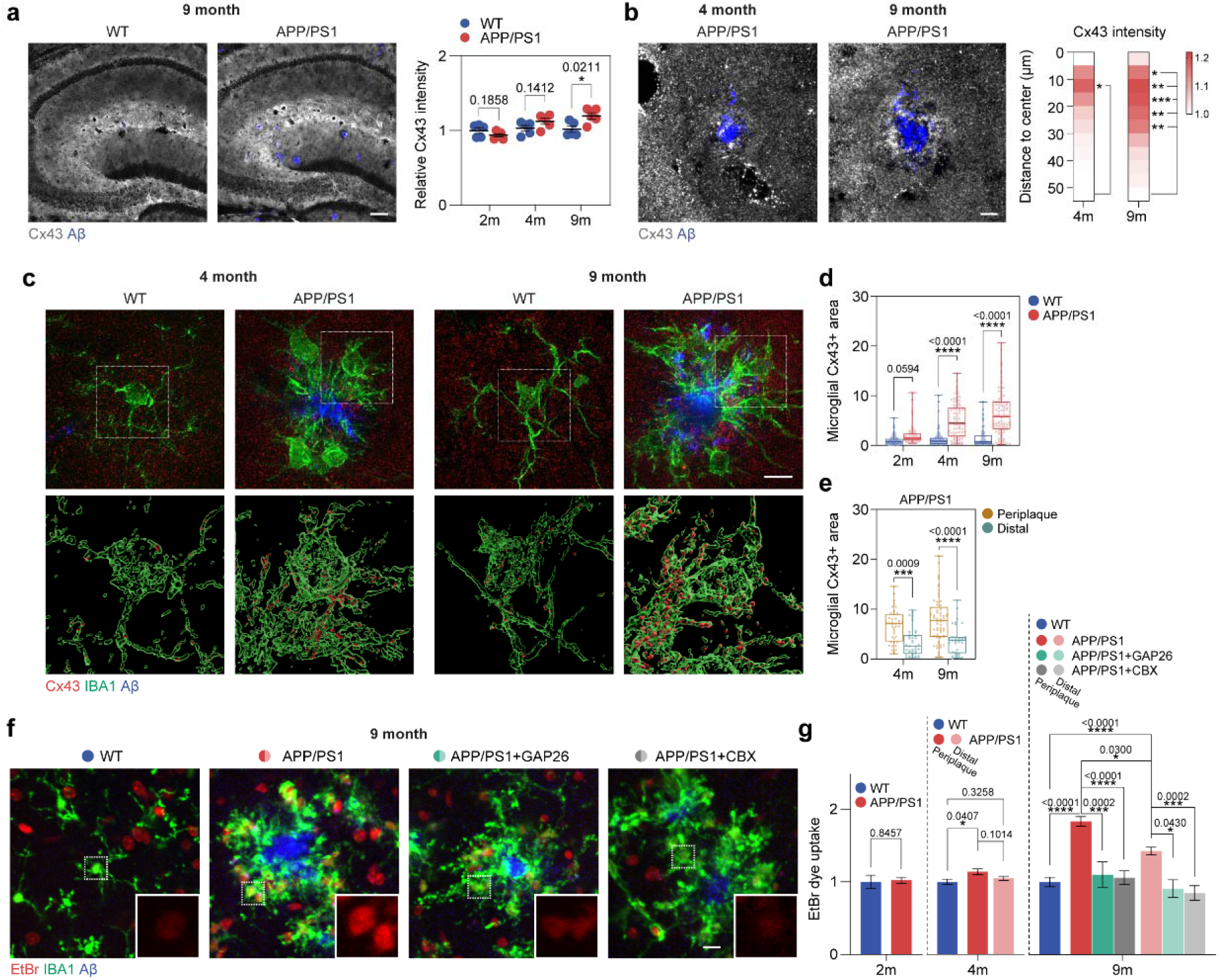
Increased microglial Cx43 correlates with Aβ pathology in AD mouse model. (A) Representative images of Cx43 and Aβ immunostaining in 9-month-old wild type (WT) and APP/PS1 mice, scale bar: 100 μm. Right: Quantification of Cx43 intensity in the hippocampus of WT and APP/PS1 mice at 2-, 4-, and 9-month-old. N = 5 mice. (B) Representative images of Cx43 and Aβ immunostaining in 4- and 9-month-old APP/PS1 mice, scale bar: 10 μm. Cx43 staining intensity at different distances from Aβ plaques (n = 23) was quantified and compared to the area distal to plaque (50-55 μm). (C) Representative super-resolution image of Cx43, IBA1, and Aβ staining in 4- and 9-month-old WT and APP/PS1 mice, accompanied by 3D reconstruction of Cx43 signal within IBA1+ volume. Scale bar: 10 μm. (D) Quantification of Cx43+ area within IBA1+ domain in WT and APP/PS1 mice, normalized to that of 2-month-old WT. N >80 cells from 5 mice. (E) Comparison of Cx43+ area within IBA1+ domain between periplaque and distal microglia. N > 40 cells from 5 mice. (F) Representative images of Ethidium bromide (EtBr) uptake by 9-month-old APP/PS1 acute brain slices and counterstained with IBA1. Cx43-specific channel blocker GAP26 and pan gap junction/hemichannel blocker CBX were added to 9-month-old APP/PS1 acute brain slices. Scale bar: 10 μm. (G) Quantification of EtBr dye intensity in dye uptake experiment on 2-, 4- and 9-month-old WT and APP/PS1 acute brain slices. N ≥ 5 experiments. Dot plots show mean ± SEM, each data point, and *p*-value. Significance: * *p* < 0.05, ** *p* < 0.01, *** *p* < 0.001, **** *p* < 0.0001.

To assess changes in microglia-specific Cx43 levels, we quantified the colocalization between Cx43 and IBA1 labeling. Cx43+ area co-localized with IBA1+ microglia was not significantly altered in APP/PS1 mice at 2 months of age, but increased at 4 and 9 months of age (by 276 ± 52% at 4 months, and by 354 ± 48% at 9 months, N > 80 cells from 5 mice, *p* < 0.0001 **Fig 2C-D**). Similarly, Cx43 levels in periplaque microglia at both 4 and 9 months of age were significantly increased **(Fig 2C-D)**. These results indicate a correlation between Cx43 levels in microglia and AD progression. We also examined the changes in Panx1 and Cx30 in microglia. While Panx1 was increased in APP/PS1 mice, the change was not age-dependent, nor it was significantly altered in periplaque microglia compared to the distal microglia (**Fig S2C**). Microglial Cx30 was not significantly changed in APP/PS1 mice compared to WT (**Fig S2D**).

Altered Cx43 function was reported in AD ^24^ and it correlated with Aβ pathology ^27^. Connexins in reactive microglia have been reported to possess both hemichannel and gap junction functions *in vitro* ^31^. However, no functional gap junction was found in surveillant or reactive microglia *in vivo* ^36^. To investigate microglial Cx43 hemichannel activity in APP/PS1 mice, we monitored ethidium bromide (EtBr) uptake in acute brain slices. At 2 months of age, no change in dye uptake by microglia from APP/PS1 mice compared to WT control was detected (**Fig 2G**). Starting from 4 months of age, we observed a significant increase in dye accumulation in the periplaque microglia (**Fig 2G**). At 9 months of age, markedly increased accumulation of EtBr was observed in both the periplaque and the distal microglia in APP/PS1 mice compared to WT mice (83.2 ± 9.2% and 42.5 ± 8.4%, N ≥ 5 experiments, *p* < 0.0001, **Fig 2F-G)**. The addition of carbenoxolone, a non-selective hemichannel blocker, normalized the dye uptake by microglia from APP/PS1 mice to the WT level **(Fig 2G)**. Because dye uptake into microglia could be mediated not only by Cx43 but also by Panx1 channels ^31^, we further used a more selective Cx43 channel blocker Gap26 ^37^, which decreased microglial dye uptake in APP/PS1 mice brain (reduced by 39.9 ± 8.3% in periplaque microglia, N ≥ 5 experiments, *p* = 0.0002), almost to the levels of WT and the CBX-treated group **(Fig 2G)**. Thus, we can attribute an increased dye uptake to an elevated Cx43 hemichannels activity in the periplaque microglia from APP/PS1 mice.

In summary, we observed a significant elevation in microglial Cx43 levels and related hemichannels activity during AD progression in APP/PS1 mice, particularly in the periplaque microglia. Our findings from AD patients and the mouse model indicate that microglial Cx43 may contribute to progressive pathogenesis of AD.

### Depletion of microglial Cx43 alleviates AD pathology and cognitive impairment

To understand the pathological role of microglial Cx43 in AD, we crossed the APP/PS1 mice with microglia-specific Cx3cr1-CreERT mice and Cx43-flox mice to achieve microglia-specific conditional Cx43 knockout in AD mouse brains (APP/PS1:Cx43^mgl-cKO^) **(Fig 3A)**. We confirmed the significant down-regulation of Cx43 in microglia in the APP/PS1:Cx43^mgl-cKO^ mice **(Fig S3A).** The changes in brain histology and cognitive performance were analyzed at 9 months of age. APP/PS1:Cx43^mgl-cKO^ mice demonstrated no changes in either the size or the number of Aβ plaques. However, Cx43 conditional knockout increased the percentage of compact plaques (plaques with a condensed core) as opposed to filamentous plaques (**Fig 3B-C**). Cx43 conditional knockout may thus reduce the neurotoxicity of the plaques, as filamentous plaques are suggested to be more neurotoxic ^38–40^.

**Fig 3.**
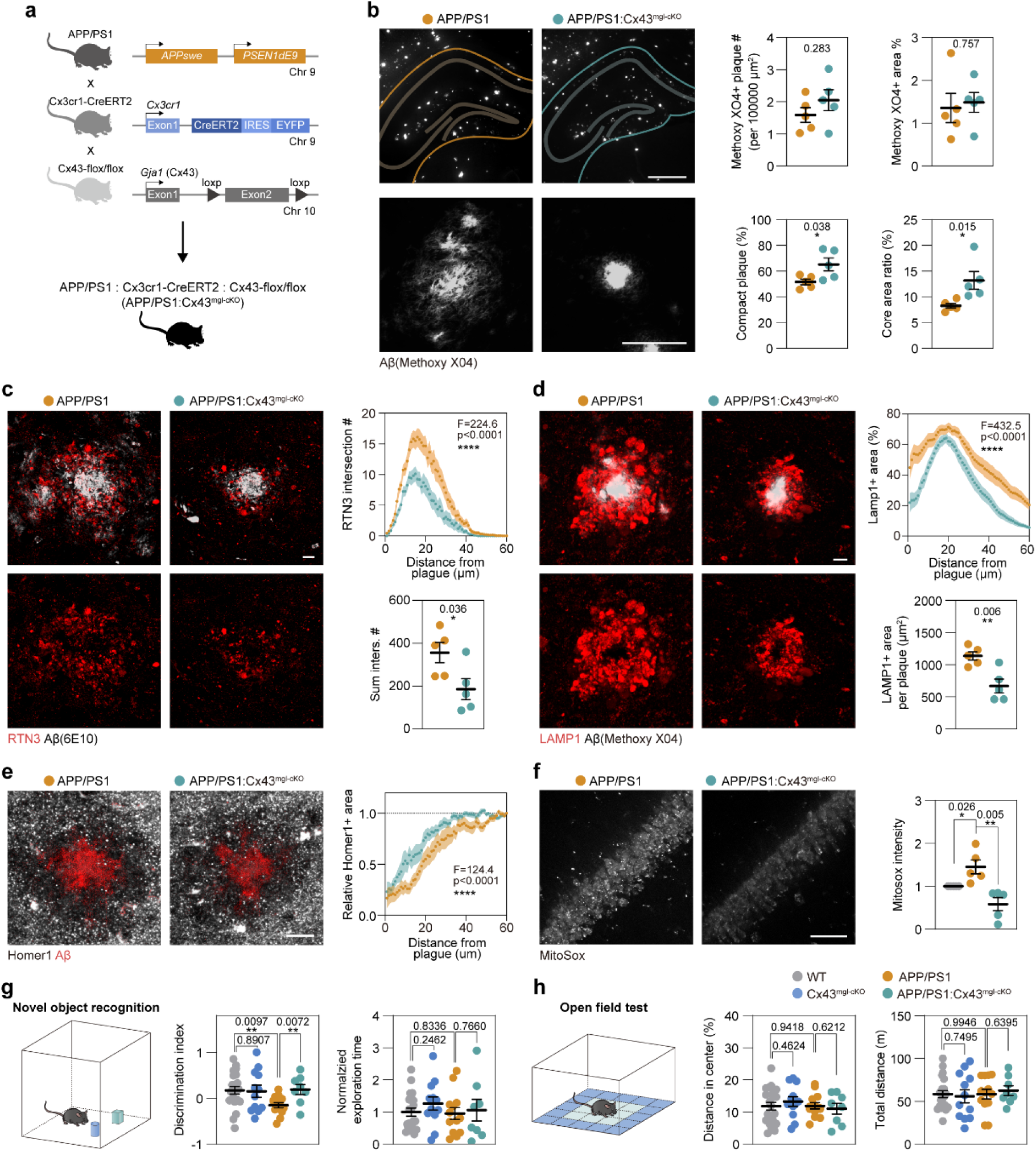
Deletion of microglial Cx43 alleviates AD pathology and cognitive impairment. (A) APP/PS1 mice were mated with Cx3cr1-CreERT and Cx43-flox/flox mice to achieve Cx43 conditional knockout in microglia (Cx43^mgl-cKO^). Non-flox or non-Cre APP/PS1 littermates were used as control. (B) Representative images of Methoxy XO4 staining in 9-month-old APP/PS1 and APP/PS1:Cx43^mgl-cKO^ mouse hippocampus. Scale bar, 50 μm. The percentage of Methoxy X04+ area and the percentage of compact Methoxy X04+ plaque were quantified. N = 5 animals. (C) Representative images of Aβ labelling by 6E10 and dystrophic neurites labelling by RTN3. Scale bar: 10 μm. The number of RTN3+ puncta at different distances from the plaque center and the sum number were quantified. N = 32 plaques. (D) Representative images of Aβ labelling by Methoxy X04 and dystrophic neurites labelling by LAMP1. Scale bar: 10 μm. LAMP1+ area % at different distances from plaque and the total area per plaque were quantified. N = 32 plaques. (E) Representative images of Aβ and postsynaptic element labelling by Homer1. Scale bar: 10 μm. Homer1+ puncta area percentage at different distances from plaque was quantified. N = 32 plaques. (F) MitoSox Red staining assay was performed on acute brain slices to analyze hippocampal CA1 neuronal oxidative stress. Scale bar, 50 μm. MitoSox intensity was quantified and normalized to the WT control of each experiment. N = 5 animals. (G) Novel object recognition test was performed on WT (N = 21), Cx43^mgl-cKO^ (N = 12), APP/PS1 (N = 14), APP/PS1:Cx43^mgl-cKO^ (N = 8) mice to examine their cognitive function, and the discrimination index was quantified. (H) Open field test was performed to examine the anxiety traits and locomotor function. Distance spent in the center zone and the total distance were quantified. Dot plots show mean ± SEM, each data point, and *p*-value. Significance: * *p* < 0.05, ** *p* < 0.01, *** *p* < 0.001, **** *p* < 0.0001.

To confirm this, we used reticulon-3 (RTN3) and lysosomal associated membrane protein 1 (LAMP1) antibodies to assess periplaque neurite dystrophy; both proteins are commonly used as dystrophic neurite markers in AD brains and animal models. LAMP1, a lysosomal marker, is highly enriched in the inner layer of periplaque dystrophic neurites ^41^, while RTN3, a tubular endoplasmic reticulum marker, is enriched in the outer layer of periplaque dystrophic neurites ^42^. APP/PS1:Cx43^mgl-cKO^ mice showed less RTN3+ or LAMP1+ dystrophic neurites surrounding the plaques compared to APP/PS1 mice (**Fig 3D-E**). We also found an increased level of the postsynaptic marker Homer1 in the periplaque area (**Fig 3F**), suggesting alleviated synaptic loss. In addition, microglial Cx43 knockout attenuated neuronal oxidative stress in APP/PS1 mice, as indicated by reduced MitoSox staining intensity in the acute brain slices, reflecting reduced superoxide production (**Fig 3G**). We further investigated whether microglial Cx43 knockout affected other manifestations of AD-related pathology. In particular, using immunostaining, we found decreased myelin loss (as assessed by MBP immunostaining) in APP/PS1 mice with microglial Cx43 ablation **(Fig S3B)**. Downregulation of microglial Cx43, however, did not affect the number or the area of GFAP+ astrocytes **(Fig S3C)**. Conditional knockout of microglial Cx43 improved cognitive performance as evidenced by results of the novel object recognition test (**Fig 3H**). No anxiety-like behaviors or locomotive defects were observed in APP/PS1:Cx43^mgl-cKO^ or APP/PS1 mice compared to WT in the open field test (**Fig 3I**). These findings suggest that ablation of Cx43 in pathological microglia mitigates neuropathology and cognitive decline during AD progression.

### Turning microglia neuroprotective by microglial-specific Cx43 knockout in AD mice

To understand the mechanisms underlying the beneficial effects of suppressing microglial Cx43 on neuropathology and cognitive impairment, we compared the cell densities and cellular morphology of hippocampal microglia in APP/PS1:Cx43^mgl-cKO^, APP/PS1, and WT mice. While APP/PS1 mice displayed a significant increase in microglia numbers compared to the WT controls, the number of microglia was even higher in APP/PS1:Cx43^mgl-cKO^ mice (**Fig 4A-B**). Increased microglial density in APP/PS1:Cx43^mgl-cKO^ mice was due to an increased population of periplaque microglia, while the number of microglia distal to plaques did not significantly differ from the APP/PS1 mice (**Fig 4B**). Arguably, periplaque microglia engage closely with the Aβ plaques to limit the damage to the adjacent neuropil ^39,40,43^. Given the increased compactness of Aβ plaques and reduced plaque-associated neural damage observed in APP/PS1:Cx43^mgl-cKO^ mice, we speculated that the microglial barrier surrounding the plaques was strengthened by Cx43 knockout. Upon closer examination of individual plaques, it became evident that APP/PS1:Cx43^mgl-cKO^ mice exhibited a higher number of periplaque microglia compared to APP/PS1 mice when normalized to the plaque perimeter or the number of plaques (**Fig 4C; Fig S4A**). Quantification of Aβ plaque volume within the microglial domain using Imaris software also showed that microglia with Cx43 knockout displayed increased interaction with Aβ plaques (**Fig 4D; Fig S4B**), which may promote the increased compactness of Aβ plaque. We also evaluated the morphology of microglia using Sholl analysis. Both periplaque and distal microglia in APP/PS1 showed reduced process complexity compared to WT microglia; Cx43 knockout increased morphological complexity of the distal microglia in APP/PS1 mice, though that of the periplaque microglia was not altered (**Fig S4C**).

**Fig 4.**
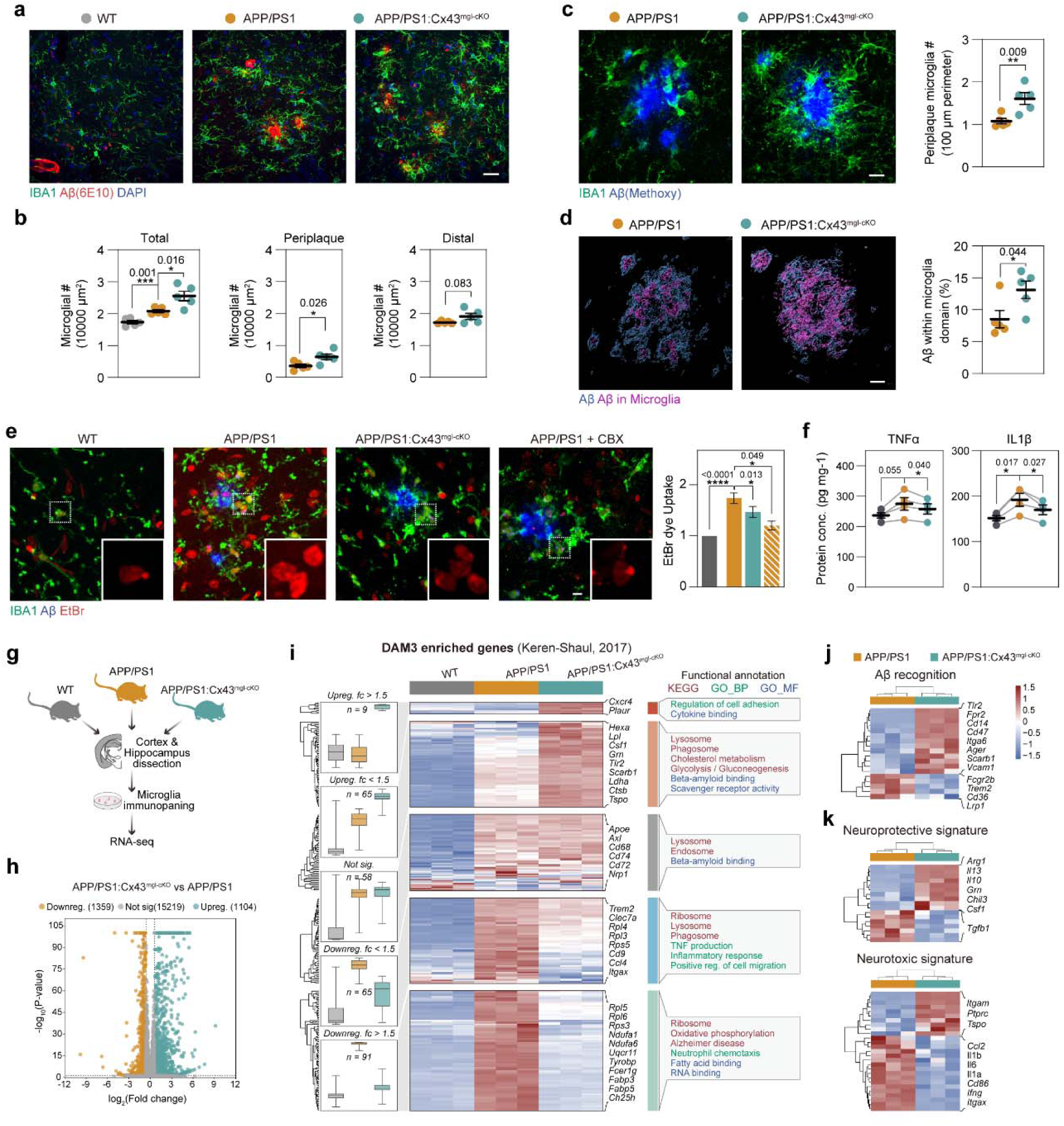
Microglia-specific Cx43 knockout induces microglia neuroprotective reactive state in APP/PS1 mice. (A) Representative image of IBA1, Aβ and DAPI staining of mouse hippocampus. (B) The total number of IBA1+ microglia, periplaque microglia, and distal microglia was quantified and normalized to the area. N = 5 animals. (C) Representative image of IBA1, Aβ and DAPI staining. For each plaque, the number of periplaque microglia was quantified, and normalized to the plaque perimeter. N = 5 animals. (D) 3D reconstruction image of panel (C) displaying Aβ plaque and the fraction of Aβ plaque within IBA1+ microglia domain. The ratio between plaque within microglia and total plaque volume was quantified. N = 5 animals. (E) Dye uptake experiment was performed on acute brain slices and counterstained with IBA1 to analyze the microglial hemichannel opening. EtBr intensity was quantified and normalized to the WT group for each experiment. N = 5 animals. (F) ELISA analysis of TNFα and IL1β cytokine level in acute brain slices normalized to protein content. (G) Bulk RNA-seq was performed on microglia isolated from WT, APP/PS1, and APP/PS1:Cx43^mgl-cKO^ mouse cortex and hippocampus. (H) Volcano plot highlighting the differentially expressed gene between APP/PS1:Cx43^mgl-cKO^ microglia and APP/PS1. (I) Heatmap shows expression of disease associated microglia 3 (DAM3) enriched genes, which were split into 5 clusters according to their expression change in APP/PS1:Cx43^mgl-cKO^ microglia compared to APP/PS1. Left: normalized gene expression level of 4 clusters. Right: functional annotation and representative genes of 4 clusters. (J-K) Heatmap shows differentially expressed genes that are associated with Aβ recognition, as well as the neuroprotective or neurotoxic signature of microglia. Dot plots show mean ± SEM, each data point, and *p*-value. Significance: * *p* < 0.05, ** *p* < 0.01, *** *p* < 0.001, **** *p* < 0.0001.

Consistent with our observation of microglial Cx43 operating as hemichannels (as shown in Fig. 2G), Cx43 knockout in microglia significantly reduced the hemichannels activity, as indicated by the reduced EtBr uptake in IBA1+ microglia in the brain slice **(Fig 4E)**. Cx43 hemichannels opening was proposed to aggravate proinflammatory response by an autocrine mechanism that activates inflammasome ^44,45^. Microglial Cx43 may play a similar role in AD pathology. Indeed, treatment with minocycline, an anti-inflammatory agent ^46^, suppressed the microglial hemichannels opening in acute brain slices from APP/PS1 mice **(Fig S4D)**, whilst knockout of microglial Cx43 significantly reduced the levels of proinflammatory cytokines TNFα and IL1β in acute brain slices **(Fig 4F)**. This suggests that an inflammatory milieu in AD facilitated the opening of the microglial Cx43 hemichannels, which may drive microgliosis to a neurotoxic pro-inflammatory phenotype.

To further understand how Cx43 ablation affects the microglial reactive landscape, we performed RNA sequencing of acutely isolated microglia from APP/PS1, APP/PS1:Cx43^mgl-cKO^, and age-matched WT control mice (**Fig 4G**). The differential expression analysis showed that Cx43 knockout in microglia promoted a distinctive neuroprotective reactive phenotype (**Fig 4H-K, Fig S4E**). Specifically, we analyzed the expression of previously reported genes enriched in the so-called disease-associated microglia (DAM) ^12^ in WT, APP/PS1, and APP/PS1:Cx43^mgl-cKO^ mice (**Fig 4I**). Our RNA-sequencing data confirmed the overexpression of DAM-associated genes in the APP/PS1 mice compared to WT mice (**Fig 4I**). We then compared the expression of these genes between APP/PS1 and APP/PS1:Cx43^mgl-cKO^ mice and identified 5 clusters according to the differential expression (**Fig 4I**). Ablation of microglial Cx43 upregulated or maintained a set of DAM genes which are associated with Aβ binding and microglia attraction to Aβ, such as *Tlr2* ^47^, *Scarb1* ^48^, *Apoe* ^49^ (**Fig 4I-J**), thus consistent with the histological observation that Cx43 knockout microglia interact more closely with the Aβ plaques. Lysosome- and phagosome-related genes were not prominently altered by Cx43 knockout (**Fig 4I**), suggesting an unaffected phagocytosis capacity.

We also observed a reduced expression of genes involved in the ribosome-, oxidative phosphorylation-, and AD-related pathways in Cx43 knockout microglia (**Fig 4I**). In addition, the downregulation of genes linked to TNF production and inflammatory-related pathways (**Fig 4I**), as well as IL1β (**Fig 4K**), are consistent with our ELISA data.

RNA-sequencing analysis also provides another aspect of how microglial Cx43 knockout-mediated phenotypic shift could ameliorate AD pathology. We observed that the neuroprotective reactive microglial signature was upregulated in APP/PS1:Cx43^mgl-cKO^ mice, including *Arg1*, *Chil3*, and *Grn*, while a majority of the neurotoxic signature genes, such as *Il1b*, *Ifng*, and *Itgax* were downregulated (**Fig 4K**). Concomitantly, immunostaining showed increased levels of ARG1 in periplaque microglia of Cx43 knockout mice compared to APP/PS1 control **(Fig S4F)**. Consistently, Cx43 knockout in microglia upregulated several genes which have been shown to be neuroprotective in AD pathology, including *Grn* ^50^ and *Tspo* ^51,52^; and several neurotoxic or pro-inflammatory genes related to AD pathology such as *Clec7a* ^53^, *Ch25h* ^54^, and *Tyrobp* ^55^ were reduced by Cx43 ablation (**Fig 4I**).

Together, these results suggest that ablation of microglial Cx43 promotes a transformation of reactive microglia towards neuroprotective phenotype in AD, establishing a closer interaction with Aβ plaques that limits the toxicity of the latter, while also reducing the expression of neurotoxic factors, which, we suggest, is due to the inhibition of Cx43 hemichannels activity.

### A lipid nanoparticle-based drug delivery strategy inhibits Cx43 hemichannels

Our finding that the neuroprotective state of microglia can be promoted by Cx43 knockout, combined with our previous observation that inhibition of astroglial Cx43 hemichannels can protect neurons in APP/PS1 mice ^24^, supports the idea that targeting Cx43 hemichannels is a promising therapeutic approach for AD treatment.

We, therefore, targeted Cx43 hemichannels using a mimetic peptide-based approach. TAT-Cx43_266-283_ peptide was designed to mimic the SH3-domain of the Cx43 C-terminus ^56^ (**Fig 5A**). Treatment with TAT-Cx43_266-283_ inhibits Cx43 hemichannels but not the gap junction channels *in vitro* ^57^. In addition, TAT-conjugated peptide readily crosses the blood-brain barrier (BBB) ^58^, supporting potential *in vivo* efficacy, which, however, has not been tested previously.

**Fig 5.**
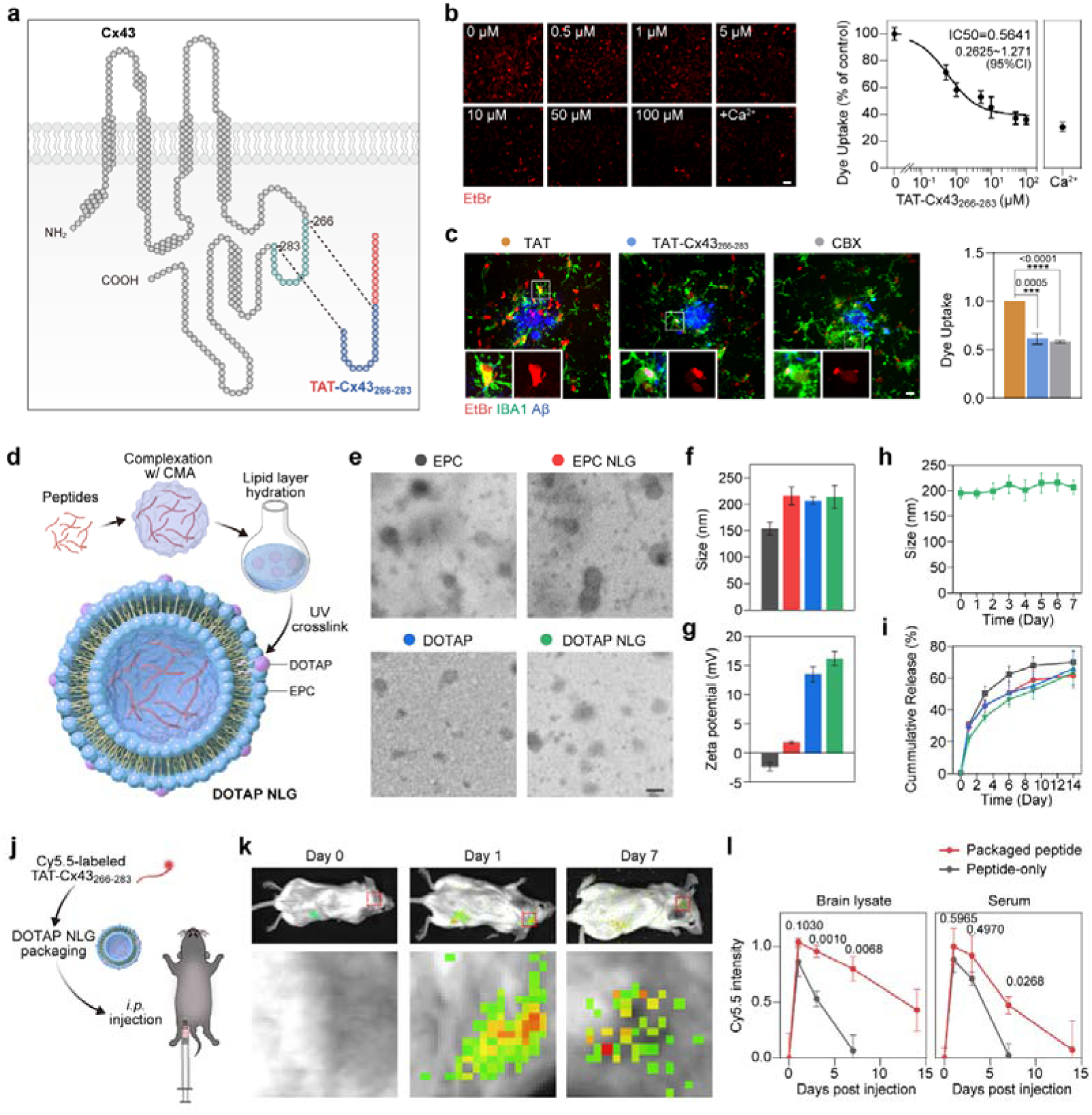
A lipid nanoparticle-based drug delivery inhibits Cx43 hemichannel activity. (A) Illustration of Cx43 structure highlighting the 266-283 region. TAT-Cx43_266-283_ is the mimetic peptide of this region conjugated to a cell-penetrating peptide TAT. (B) Dye uptake experiment was performed on primary microglia in combination with TAT-Cx43_266-283_ treatment at different concentrations. The addition of 1.8 mM Ca^2+^ which inhibits hemichannel opening ^59^ serves as negative control. Scale bar, 10 μm. EtBr signal was quantified, and a non-linear regression analysis was performed to determine the IC50 value. N = 3 coverslips. (C) 50 μM TAT-Cx43_266-283_ peptide was added to acute brain slices for 15 min before the EtBr uptake experiment. Followed by IBA1 counter-immunostaining. EtBr signal intensity in IBA1+ microglia was quantified and normalized to TAT-treated APP/PS1 control. N = 4 experiments. (D) Schematic diagram of TAT-Cx43@LNP construction. (E) Electronic microscopic of lipid nanoparticles with different compositions. (F) Measurement of the size of lipid nanoparticles. (G) Measurement of zeta potential of lipid nanoparticles. (H) The size of lipid nanoparticles at different days post-production. (I) Measurement of the peptide release rate at 37 □ of lipid nanoparticles *in vitro*. (J) Cy5.5-labeled TAT-Cx43_266-283_ peptide packaged with DOTAP nanolipid gel (DOTAP-NLG) was i.p. injected into mice, followed by biofluorescence imaging at different time points. (K) representative images of biofluorescence imaging at days 0, 1, and 7 post-injection, revealing the retainment of peptides in the brain. (L) Microplate reader analysis of Cy5.5 intensity of brain lysate and serum collected at day 0, 1, 3, 7, 14 post-injection of TAT-Cx43 peptide alone or with DOTAP-NLG packaging. N = 3 (peptide-alone) or 5 (with packaging) experiments. Dot plots show mean ± SEM, each data point, and *p*-value. Significance: * *p* < 0.05, ** *p* < 0.01, *** *p* < 0.001, **** *p* < 0.0001.

We first confirmed the capability of the TAT-Cx43_266-283_ to inhibit Cx43 hemichannels. In the primary microglial culture, TAT-Cx43_266-283_ effectively suppressed hemichannels opening as assessed by the EtBr uptake assay (IC_50_ = 0.5641 μM, 95% confidence interval = 0.2625∼1.271 μM, n = 3 experiments) (**Fig 5B**). In acute brain slices from APP/PS1 mice, treatment with 50 μM of TAT-Cx43_266-283_ was sufficient to inhibit microglial Cx43 hemichannels (assessed by EtBr uptake) with an efficacy similar to that of 200 μM CBX; both reduced hemichannels activation to ∼60% of the control level (**Fig 5C**). In primary microglia treated with Aβ_25-35_, exposure to 50 μM of TAT-

Cx43_266-283_ modified the microglial reactive state, as indicated by the increased ARG1 and reduced iNOS staining (**Fig S5A-B**). Furthermore, as Cx43 is also expressed in astrocytes forming both hemichannels and gap junction channels ^59^, we examined the effect of TAT-CX43_266-283_ on astrocytes. In acute brain slices, TAT-Cx43_266-283_ inhibited astrocytic Cx43 hemichannels opening (shown by EtBr uptake assay), but did not affect gap junctions as determined by gap-FRAP assay, which reveals the extent of syncytial coupling (**Fig S5C-D**). These data support the notion that TAT-Cx43_266-283_ can effectively and specifically inhibit Cx43 hemichannels and modulate microglial reactivity without affecting Cx43 gap junction functions.

However, peptide proteins normally have a short half-life *in vivo*, which limits their bioavailability ^60^. To address this limitation, we adopted a liposome-based peptide drug delivery system. We packaged the peptide with the 1,2-dioleoyl-3-trimethylammonium propane (DOTAP)-based, chitosan-methacrylate (CMA)-crosslinked, nanolipid gel (**Fig 5D**), initially designed for siRNA delivery ^61^. We named the packaged TAT-Cx43_266-283_ formulation ‘TAT-Cx43@LNPs’. Electron microscopy revealed that the addition of DOTAP and/or CMA crosslinking slightly increased the size of the nanolipid particle compared to the basic egg phosphatidylcholine (EPC) particle, while the addition of DOTAP significantly increased the surface potential of the nanolipid particles. (**Fig 5E-G**). The size of the nanolipid gel remained stable *in vitro* for 7 days, and could achieve sustained, stable release of peptides for 14 days (**Fig 5H-I**). Next, we examined the capacity of the sustained release of the lipid nanoparticle *in vivo*. The peptide was labeled with fluorophore Cy5.5 before being packaged with lipid nanoparticles and was subsequently intraperitoneally injected into mice (**Fig 5J**). Biofluorescence imaging detected Cy5.5 signal in the brain for up to 7 days post injection (**Fig 5K**). To further confirm the tissue retainment of the peptide drug, we isolated tissues post-injection and homogenized them to analyze the fluorescence intensity. When peptides were injected without LNP packaging, they peaked on 1-day post-injection but were absent on 7^th^ day post-injection in the brain lysate and serum (**Fig 5I**). However, when LNP-packaged, peptides display significantly longer stability *in vivo*. On 14^th^ day post-injection, LNP-packaged peptides were maintained at a relatively high level in the brain lysates (42.9 ± 17.1% compared to day 1, N = 6 mice, *p* = 0.0068) but were not detectable in serum (**Fig 5I**). Our LNP packaging method moderately elevated the peptide level in the brain, but could not achieve exclusive brain delivery, as Cy5.5-peptides were also found in the heart, liver, and kidney (**Fig S5E**). Nonetheless, these data show that TAT-Cx43@LNPs can achieve sustained drug release and brain retention, substantiating their feasibility for *in vivo* application.

We further examined the biosafety of the TAT-Cx43@LNPs. Injection of TAT@LNPs or TAT-Cx43@LNPs did not significantly change the mouse weight upon consecutive weekly injections for 6 weeks (**Fig S5F**). Cx43 is highly expressed in several vital organs, especially in the heart, where Cx43 contributes to action potential propagation ^62^. To investigate potential adverse effects on the cardiac function, we performed electrocardiograms on mice before and after the treatment. We found that TAT-Cx43@LNPs altered neither the heart rate nor the propagation of electrical signal across the heart, as judged by the duration of P wave, P-R interval, QRS complex, and Q-T interval, compared to the baseline measurement or the TAT@LNP treated control mice (**Fig S5G-H**). We also performed the serum biochemistry test before and after treatment to examine the impact on vital organ functions, including the heart, liver, kidney, muscle, and pancreas. It appeared that LNP treatment did not significantly affect serum markers of organ functions (**Fig S5I**). In summary, our nanocarrier-based TAT-Cx43_266-283_ delivery strategy appears to be safe without apparent off-target effects.

### Nanocarrier-delivered Cx43 blocker rescues neuropathology and cognitive decline in AD mice

To test whether TAT-Cx43@LNPs can affect AD progression, we treated WT and APP/PS1 mice with TAT-Cx43@LNPs once per week (120 μg/g, peptide 8.2 nmol/g, i.p.) starting from 8 months of age. After 6 weeks of treatment, mice were subjected to behavioral tests and histological analysis **(Fig 6A)**. The TAT@LNPs treated mice acted as control (which did not affect cognitive outcome as indicated by the novel object test; **Fig S6A**). In the Barnes Maze test, TAT-Cx43@LNPs treated APP/PS1 mice spent more time in the target quadrant in the probe trial compared to the TAT@LNPs treated APP/PS1 mice **(Fig 6B-C; Fig S6B-C)**, suggesting an improved spatial memory. In the novel object recognition test, TAT-Cx43@LNPs treated APP/PS1 mice showed a higher preference for the new object compared to the TAT-treated APP/PS1 mice, indicating an improved short-term memory **(Fig 6D)**. However, TAT-Cx43@LNPs treatment did not affect the locomotive activity or anxiety-like behavior in the open field test **(Fig 6E; Fig S6D)**. TAT-Cx43@LNPs treatment in the WT mice affected neither cognitive function, nor locomotive activity, nor anxiety-like behavior **(Fig 6B-E; Fig S6B-D)**. In summary, these results indicate that TAT-Cx43@LNPs restored the cognitive functions of APP/PS1 mice.

**Fig 6.**
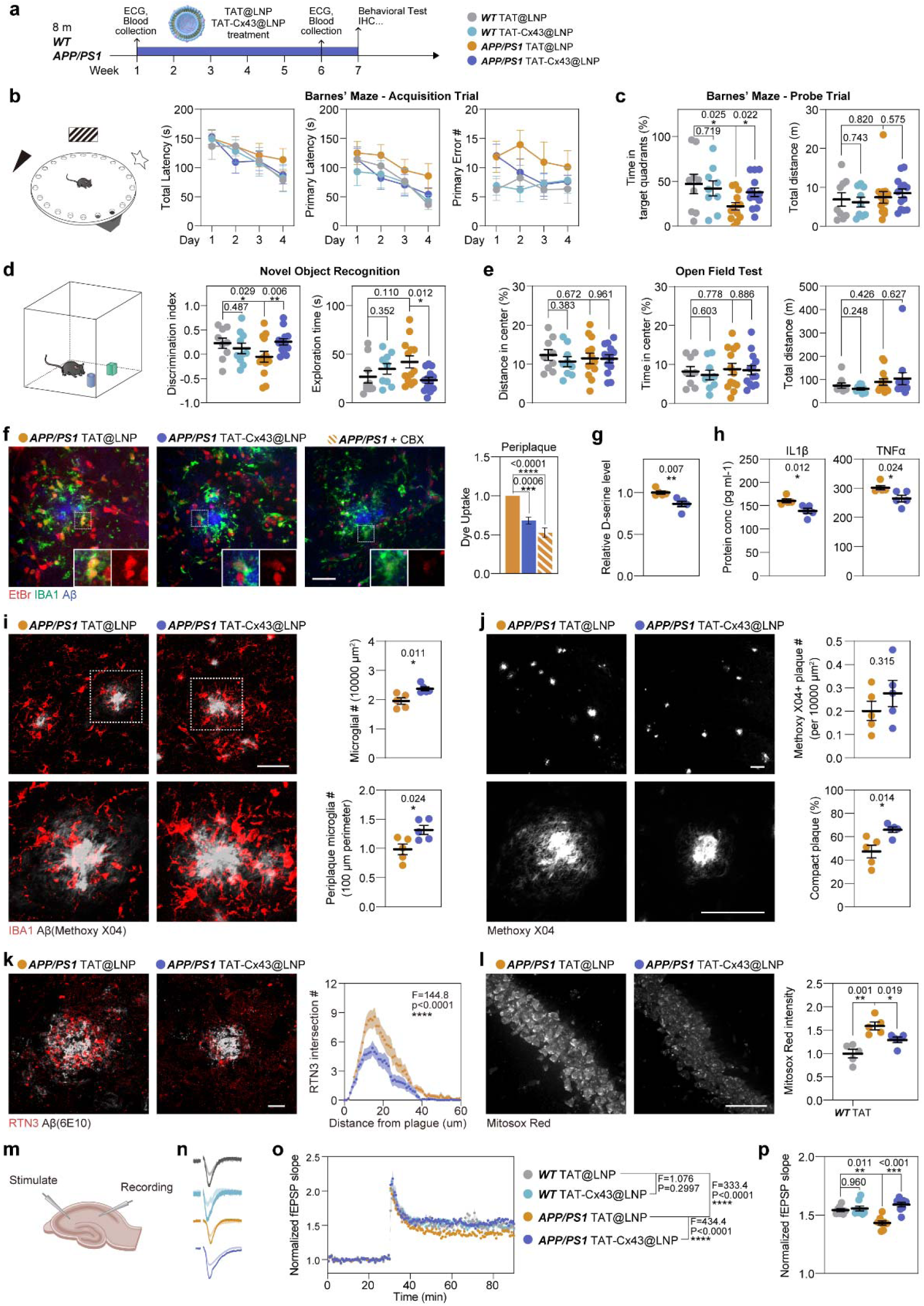
Nanolipid-delivered Cx43 blocker rescues neuropathy and cognitive decline in AD mouse model. (A) 8-month-old WT and APP/PS1 mice were treated with TAT-Cx43@LNP or with TAT@LNP as control weekly for 6 weeks, before being subjected to behavioral tests and further histology analysis. (B) Barnes’ maze test was used to analyze mouse cognitive function, specifically, the spatial learning and memory function. The total latency to the target hole, primary latency, and primary exploratory error during the acquisition phase were quantified. (C) The time spent in the target quadrant and the total distance during the probe trial. For WT TAT, WT TAT-CX43, APP/PS1 TAT, and APP/PS1 TAT-CX43, N = 9, 9, 13, 13 animals. (D) Novel object recognition test was used to examine the cognitive function. The discrimination index and the total object exploration time were quantified. (E) Open field test was performed to examine the anxiety traits and locomotor function. Distance and time spent in the center zone and the total distance were quantified. (F) Dye uptake experiment was performed on acute brain slices followed by IBA1 counterstaining. EtBr signal intensity in IBA1+ microglia was quantified and normalized to TAT-treated APP/PS1 control. Scale bar, 50 μm. N = 5 animals. (G) D-serine level in the supernatant of acute brain slices was measured and normalized to the TAT@LNP treated APP/PS1 group. N = 5 animals. (H) Proinflammatory cytokine IL1β and TNFα levels in acute brain slices measured by ELISA. N = 5 animals. (I) Representative images of IBA1 and Methoxy X04 staining. Scale bar, 50 μm. The number of microglia in hippocampus and the periplaque microglia number were quantified. N = 5 animals. (J) Representative images of Methoxy X04 staining in the hippocampus. Scale bar, 50 μm. The number of Aβ plaques number and the percentage of compact plaques were quantified. N = 5 animals. (K) Representative images of RTN3 and Aβ staining in the hippocampus. Scale bar, 10 μm. RTN3+ puncta number at different distances from plaque was quantified. N = 45 (TAT) and 42 (TAT-Cx43) plaques. (L) MitoSox Red staining assay was performed on acute brain slices to analyze hippocampal CA1 neuronal oxidative stress. Scale bar, 50 μm. MitoSox intensity was quantified and normalized to the WT control. N = 5 animals. (M) Theta burst stimulation-induced long-term potentiation (TBS-LTP) was measured on acute brain slices to examine hippocampal synaptic plasticity. (N) Representative field excitatory post synaptic potential (fEPSP) recording before and after LTP induction. (O) Quantification of fEPSP slope during LTP recording experiment. (P) Normalized fEPSP slope after LTP induction. Dot plots show mean ± SEM, each data point, and *p*-value. Significance: * *p* < 0.05, ** *p* < 0.01, *** *p* < 0.001, **** *p* < 0.0001.

TAT-Cx43@LNPs suppressed the Cx43 hemichannels in both microglia **(Fig 6F)** and astrocytes **(Fig S6G)** as determined by the dye uptake assay in the acute brain slices, but did not affect Cx43 protein levels **(Fig S6E-F)** or gap junctional communication (as judged by dye coupling) in the astrocytes **(Fig S6H)**. Furthermore, TAT-Cx43@LNPs reduced the extracellular level of D-serine **(Fig 6G)**. TAT-Cx43@LNPs also made microglia less pro-inflammatory, as reflected by the reduced production of proinflammatory cytokine TNF-α and IL1-β **(Fig 6H)**, as well as an increased ARG1 expression **(Fig S6I)**. Similarly to the APP/PS1:Cx43^mgl-cKO^ mice, TAT-Cx43@LNPs treatment increased microglial numbers, especially of the periplaque microglial population (**Fig 6I; Fig S6J-K**), increased microglia-plaque interaction (**Fig S6L**), and slightly increased microglial ramification (**Fig S6M**). These observations are consistent with those in the APP/PS1:Cx43^mgl-cKO^ mice, supporting the notion that the inhibition of microglia Cx43 hemichannels promotes a neuroprotective microglia reactive state.

We next examined whether TAT-Cx43@LNPs rescued the neuropathology in APP/PS1 mice. Immunohistology showed that while TAT-Cx43@LNPs did not significantly alter the total area or the number of Aβ plaques, instead, it increased their compactness (**Fig 6J, Fig S6N**), which was accompanied by reduced plaque-associated neurite dystrophy (**Fig 6K**). TAT-Cx43@LNPs treatment mitigated the neuronal oxidative stress reflected by MitoSox staining of brain slices (**Fig 6L**). In addition, TAT-Cx43@LNPs treatment ameliorated the myelin loss, as reflected by the elevated MBP+ area **(Fig S6O)**. Finally, TAT-Cx43@LNPs treatment rescued the reduced long-term potentiation (LTP) in APP/PS1 mice. (**Fig 6M-P**).

Collectively, our findings demonstrate that TAT-Cx43@LNPs can be delivered into the CNS to effectively inhibit microglial Cx43 hemichannels, mitigate neuropathology, and restore impaired neuronal functions and cognitive deficits.

### TAT-Cx43@LNPs prevent neuropathology at the early stage of AD

We demonstrated that ablation of microglial Cx43 or pharmacological suppression of Cx43 hemichannels did not alter the plaque load. It has been shown, however, that microglia differentially affect the plaque load at different stages of AD pathology ^40^. To investigate whether suppression of Cx43 hemichannels by TAT-Cx43@LNPs at the early stage of AD can arrest or delay AD pathology, we treated APP/PS1 mice with TAT-Cx43@LNPs weekly from 3.5 to 5.5 months of age **(Fig 7A)**. We first verified that TAT-Cx43@LNPs treatment effectively inhibited Cx43 hemichannels in both microglia and astrocytes in APP/PS1 mice at 5.5 months using the EtBr uptake experiment **(Fig S7A)**. MitoSox staining of the acute brain slices revealed that TAT-Cx43@LNPs reduced the oxidative stress in hippocampal neurons compared to the LNP-TAT treated control **(Fig 7B)**. Finally, TAT-Cx43@LNPs alleviated neurite dystrophy as revealed by a reduced RTN3 staining and increased Homer1 staining intensities **(Fig 7C-D)**, suggesting that administration of TAT-Cx43@LNPs at early ages can delay the neuropathological progression in APP/PS1 mice.

**Fig 7.**
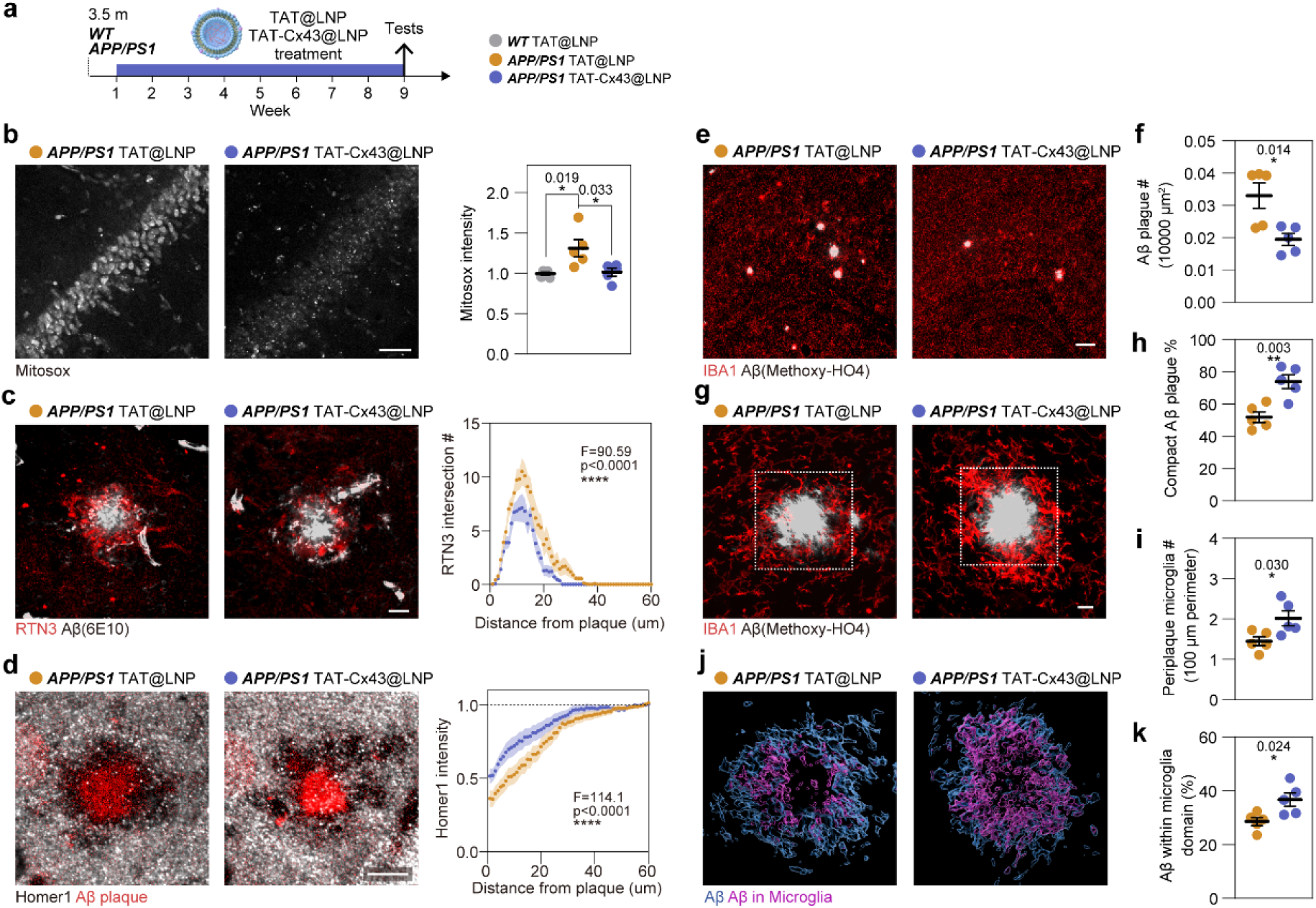
TAT-Cx43@LNPs prevent neuropathogenesis at the early stage of AD pathogenesis. (A) 3.5-month-old WT and APP/PS1 mice were treated with TAT-Cx43@LNP or with TAT@LNP as control weekly for 8 weeks, before being subjected to histology analysis. (B) MitoSox Red staining assay was performed on acute brain slices to analyze hippocampal CA1 neuronal oxidative stress. Scale bar, 50 μm. MitoSox intensity was quantified and normalized to the WT control. N = 5 animals. (C) Representative images of RTN3 and Aβ staining in the hippocampus. Scale bar, 10 μm. RTN3+ puncta number at different distances from plaque was quantified. N = 26 (TAT) and 29 (TAT-Cx43) plaques. (D) Representative images of Aβ and postsynaptic element labelling by Homer1. Scale bar: 10 μm. Homer1+ puncta intensity at different distances from plaque was quantified. N = 11 (TAT) and 13 (TAT-Cx43) plaques. (E) Representative images of IBA1 and Methoxy X04 staining. Scale bar, 50 μm. (F) Quantification of Aβ plaque number. N = 5 animals. (G) Higher resolution images of IBA1 and Methoxy X04 staining. Scale bar, 10 μm. (H) Quantification of the percentage of compact plaque. N = 5 animals. (I) For each plaque, the number of periplaque microglia was quantified, and normalized to the plaque perimeter. N = 5 animals. (J) 3D reconstruction image of the panel (G) displaying Aβ plaque and the fraction of Aβ plaque within IBA1+ microglia domain. (K) The ratio between plaque within microglia and total plaque volume was quantified. N = 5 animals. Dot plots show mean ± SEM, each data point, and *p*-value. Significance: * *p* < 0.05, ** *p* < 0.01, *** *p* < 0.001, **** *p* < 0.0001.

In contrast to the treatment at the late stage of AD, early TAT-Cx43@LNPs treatment significantly reduced the number of Aβ plaques as shown by the Methoxy-X04 staining (**Fig 7E-F**). Early TAT-Cx43@LNPs treatment also increased the compactness of Aβ plaques, similar to the late-stage treatment (**Fig 7G-H, Fig S7C**). Alleviated Aβ pathology by the TAT-Cx43@LNPs treatment was associated with the enhanced microglia-plaque interactions, as shown by the increased number of periplaque microglia, as well as the microglia-interacting fraction per plaque (**Fig 7I-K**). Additionally, TAT-Cx43@LNPs increased the ramification of microglia distal to plaques (**Fig S7D**). Our findings highlight the potential therapeutic value of suppressing Cx43 hemichannel activity as a novel approach for both the prevention and treatment of AD.

## Discussion

This study pioneers a cell-targeted treatment strategy for AD, based on switching microglia to neuroprotective phenotype by targeting microglial Cx43 hemichannels; to the best of our knowledge this is the first report of microglia-targeting therapy. We provided comprehensive evidence to support the role of microglial Cx43 hemichannels in the pathogenesis of AD in patients and in a mouse model of AD. We found an increased presence of microglial Cx43 in both AD patient brains and in APP/PS1 mice, suggesting the relevance to human disease. We subsequently demonstrated that microglial Cx43 operates as hemichannel and, for the first time, provided *in vivo* evidence to show that inhibition of microglial Cx43 hemichannels alleviates AD progression by inducing a neuroprotective microglial state. The neuroprotective microglia established a closer interaction with and promoted the compactness of the Aβ plaque, as well as reduced the expression of neurotoxic factors. We further provided preclinical evidence that targeting Cx43 hemichannels using our novel formulation TAT-Cx43@LNPs represents a promising strategy for AD management through shifting reactive microglia towards a neuroprotective state, leading to (i) more compacted Aβ plaque morphology and reduced plaque number if treated at the early stage; (ii) reduced periplaque neurite dystrophy and synaptic loss; (iii) decreased neuronal oxidative stress; (iv) rescued myelin loss; (v) ameliorated deficient LTP deficit; and last but not least, (vi) improved cognitive function. Our discoveries highlight the potential of microglial Cx43 as a therapeutic target and, importantly, set a precedent for developing cell-specific treatments that could transform AD therapy.

Although the role of microglial Cx43 in neuropathology has been suggested by *in vitro* studies, there is a lack of *in vivo* evidence; similarly, the impact of Cx43 on microglia reactive status remained unexplored. In the healthy brain, microglia express low levels of Cx43, which, however, is increased by lipopolysaccharide (LPS) or proinflammatory factors ^63–65^. Increased microglial Cx43 was also found *in vivo* around stab wounds ^66^. The suggested gap junction function of microglia connexins was only based on *in vitro* observations and was never confirmed *in vivo* ^36^. Treatment of primary cultured microglia with Aβ opens Cx43 hemichannels, triggering the release of glutamate and ATP, and inducing excitotoxicity ^27,67,68^. These observations consolidated the notion of changes in Cx43 expression and function associated with reactive microgliosis. In this study, we revealed how Cx43 hemichannels modulate reactive microgliosis in the context of AD. We observed a significant increase in microglial Cx43 hemichannels activity, especially in the periplaque microglia. We provided new evidence showing that conditional knockout of microglial Cx43 significantly reduces hemichannels activity, alters microglial reactivity, increases periplaque microglia population, and consequently alleviates neuropathology and cognitive impairments. Moreover, manipulating microglial Cx43 leads to marked transcriptomic changes in microglia in APP/PS1 mice that are instrumental in promoting neuroprotective reactive status. Further research is needed to reveal the molecular mechanism by which Cx43 hemichannels regulate gene expression.

To date, most of the AD drug-development is focused on disease-modifying therapies aimed at Aβ ^5,69^. The β amyloid cascade hypothesis, however, is insufficient to explain many facets of AD progression, leading to a 98% failure rate of clinical trials ^3,4^; hitherto, no disease-modifying therapy exists ^5^. Microglia-targeting AD therapy has gained increasing attention as a growing body of evidence indicates that microglia engage in the pathological progression of AD ^70^. However, previous attempts at suppressing reactive microgliosis using minocycline have proven ineffective in clinical trials ^71^, suggesting the need for new strategies to modulate microglia state. Our study illustrates the critical role of microglial Cx43 hemichannels in regulating the microglia response to Aβ pathology. Moreover, genetic deletion or pharmacological inhibition of microglial Cx43 alleviates AD progression. In CNS, Cx43 is highly expressed by astrocytes, forming gap junctions that mediate cell-to-cell communication within the CNS. In our previous studies, we demonstrated that Cx43 hemichannels in astrocytes disrupt the microenvironment homeostasis and contribute to AD progression ^24^; we also reported that deletion of astroglial Cx43 in APP/PS1 blocked hemichannels opening, leading to reduced neuronal damage in the hippocampus ^24^. Together, these results highlight the potential of targeting Cx43 hemichannels as a preventive and disease-modifying strategy of AD, suggesting a promising avenue for therapeutic intervention.

However, there are two major obstacles to be circumvented in targeting Cx43 hemichannels for AD treatment. Firstly, given that appropriate Cx43 gap junctions and hemichannels are essential for the function of multiple organs, any therapeutic strategy targeting Cx43 needs to be carefully controlled to minimize side effects. Our previous studies have shown that an alkaloid compound Boldine that can cross blood brain barrier can modify the neuropathology of AD by blocking Cx43 hemichannels ^72^. Boldine has recently been under clinical trial for treating overactive bladder in women, ^72^. However, its off-target effects remain unknown. There are other Cx43-targeting agents in clinical trials or preclinical tests ^73^; among them, mefloquine, a potent Cx channel blocker, combined with donepezil (THN201), is under preclinical testing for cognitive deficit linked to AD ^73^. However, mefloquine has a higher efficiency in targeting Cx36 compared with Cx43. Several other Cx43 channel blockers were used in various animal models. For instance, Gap26 and Gap27 are Cx43 mimetic peptides that target the extracellular loop of Cx43 ^74^, which can significantly impair cell-cell communications. This is because they not only block Cx43 hemichannels but also interfere with gap junctions ^75^, as well as with other connexin channels ^74^. D4 is a newly developed connexin channel blocker that showed high efficacy in suppressing reactive gliosis in an epilepsy model ^76^, but it does not discriminate between Cx26, Cx30, Cx43, and Cx45 hemichannels, which may lead to significant side effects.

TAT-Cx43_266-283_ peptide specifically inhibits the Cx43 hemichannels in astrocytes ^57^. Here, we demonstrated that TAT-Cx43_266-283_ peptide blocks Cx43 hemichannels in both microglia and astrocytes without measurable effects on astrocytic gap junctions. The safety profile of TAT-Cx43_266-283_ peptide in our preclinical study also supports its clinical translation, showing that it does not interfere with the physiological function of vital organs expressing Cx43, such as the heart, liver, and kidney. Secondly, peptide drugs have a short half-life in serum due to proteolytic digestion ^60^, an obstacle that needs to be addressed for the management of chronic diseases such as AD. To this end, we developed a novel lipid nanoparticle packaging system to formulate the TAT-Cx43@LNPs drug, which improves the bioavailability of TAT-Cx43_266-283_ peptide in the CNS, achieves long-term glial Cx43 hemichannels inhibition, and more importantly, not only prevents and rescues the Aβ-related neuropathology but also alleviates the cognitive decline. One limitation of our lipid nanoparticle packaging system is the lack of brain-specific targeting, which may lower the drug availability in the brain and increase the risk of side effects, which awaits further iteration to introduce the active brain targeting strategy ^77^. Nonetheless, our data demonstrate that targeting Cx43 hemichannels by a lipid nanoparticle-based mimetic peptide delivery has promising clinical value for AD therapy and prevention.

In conclusion, our study provides insight into the early cellular changes in AD pathology and introduce a pioneering cell-specific strategy with significant therapeutic potential. This innovative approach holds the promise of transforming AD treatment and paving the way for a new era in AD containment. Future work is needed to fully realize the treatment potential of our strategies for patients with AD in clinical trials.

## Materials and Methods

### Human samples

Human post-mortem brain tissues were provided by Human Brain Bank, Chinese Academy of Medical Sciences & Peking Union Medical College, Beijing, China, under the ethical approval of the Ethics Committee of the Chinese Academy of Medical Sciences (NO. 009-2014). After deparaffinization, the paraffin-embedded sections were treated with 3% H_2_O_2_ and processed for citric acid antigen retrieval. The sections were blocked with 2% BSA in TBST and incubated with primary antibody for 2 h at 37 □ and then overnight at 4 □, followed by a serial rinsing and incubation with fluorophore-conjugated secondary antibody. The detailed information of healthy controls and AD patients was listed in **Fig. S1B**. Primary antibodies used are Rabbit-anti-Cx43 (Merck, C6219), Rabbit-anti-PANX1 (Thermo, 487900), Goat-anti-IBA1 (Abcam, ab5076), Rat-anti-GFAP (Abcam, ab279291).

### Animals

Wild type (WT) or APP_swe_/PS1_dE9_ (APP/PS1) mice with a C57BL/6j background were obtained from GemPharmatech Co., Ltd. Cx3cr1-CreERT mice, as previously reported, were obtained from the Jackson Lab (Strain # 21160). They were mated with Cx43-flox (Strain # 008039) mice and APP/PS1 mice to generate APP/PS1: Cx3cr1-CreERT: Cx43-floxed mice (APP/PS1:Cx43^mgl-cKO^). It is worth noting that APP/PS1 transgene and Cx3cr1-CreERT knockin/knockout are both located at chromosome 9. In this study, APP/PS1 heterozygote littermates were used as a control for behavioral tests, histological analysis, acute brain slices experiments, and RNA-sequencing experiments. To induce Cx43 knockout, 30 mg/mL tamoxifen in 1:9 ethanol/corn oil mixture was given to mice via gavage at 50 μL each day for 5 consecutive days. The induction started at 8.5-month-old, and the behavioral test at 9.5-month-old. For TAT-Cx43@LNP treatment, mice received TAT@LNP or TAT-CX43@LNP (15 mg/mL, 8 μL/g) injection once per week (i.p., equivalent to 120 μg LNPs/g, and 8.2 nmol peptides/g). All animals used in the study were maintained in a PC2 pathogen-free animal facility. All procedures were performed under the ethical approval of the Sun Yat-Sen University Institutional Animal Care and Use Committee (NO. SYSU-IACUC-2024-000452).

### Behavioral tests

Before being subjected to behavioral tests, mice were handled for 3 consecutive days until they appeared calm on the handler’s hand without displaying stressful behaviors, such as jumping, biting, defecating, or urinating. Tests were conducted in the following sequence: open field test, novel object recognition, and Barnes’ maze test for the TAT-CX43@LNP experiment. Videotaping and data analysis were conducted using the VisuTrack software and hardware (Xinrun, Shanghai).

**Open field test** was used to determine the anxiety trait and locomotor function. Mice were placed in the center of a 50 × 50 × 40 (height) cm^3^ white box and were allowed to explore for 20 min. The floor was divided into 5 x 5 squares and the center 9 squares were considered as center zone.

**Novel object recognition test** was used to determine the cognitive function. In the familiarization session, mice were placed in a 25 × 25 × 40 (height) cm^3^ white box with 2 rectangular plastic objects placed 8 cm away from the walls and allowed to explore for 5 min. The test session took place two hours later, where one of the rectangular objects was replaced by a cylinder (the novel object) before the mouse was re-introduced into the apparatus. The discrimination index calculated as (Time^new^-Time^old^) / (Time^new^-Time^old^) was used to determine the exploratory preference toward the novel object.

**Barnes’ Maze test** was used to determine spatial memory and cognitive functions and performed according to a previous publication ^78^ with minor adjustments. The platform was a 92 cm diameter round plastic board with 20 equally spaced 5 cm diameter holes at the border, with one hole that could be attached to the escape tunnel. The platform was held 80 cm above the ground. Three distal visual cues were placed surrounding the platform. The experiment room was well-lit at ∼1000 lux. In the habituation phase (Day 1), the mouse was placed in the escape tunnel for 1 min and allowed to explore the platform for 5 min or until the mouse entered the tunnel. In the acquisition trial, mice were placed onto the platform with a transfer box and allowed to explore for 3 min or till the mice entered the tunnel. This procedure was conveyed twice daily with a 2-hour interval, for a total of 4 days (Day 1-4). In the probe trial (Day 5), the escape tunnel was removed, and the mouse was allowed to explore the platform for 3 min.

### Immunostaining

Mice were anesthetized by the i.p. injection of 20% urethane at 10 μL/g, followed by transcardial perfusion of PBS and 4% paraformaldehyde (PFA). The brains were then isolated and subjected to post-fixation with 4% PFA at 4 □ overnight, then immersed in 30% Sucrose for cryoprotection, before cryostat sectioning for 20 μm thick slices. Brain slices were permeabilized with 0.5% Triton-X100 in PBS (0.5% PBST) for 1 hour, blocked with 2% BSA in PBST for 1 hour, and incubated with primary antibody overnight at 4 □, followed by secondary antibody, counterstained with DAPI, and mounted for microscopic observation. Primary antibodies used are Rabbit-anti-Cx43 (Merck, C6219), Rabbit-anti-PANX1 (Thermo, 487900), Rabbit-anti-Cx30 (Thermo, 712200), Goat-anti-IBA1 (Abcam, ab5076), Goat-anti-GFAP (Abcam, ab53554), Rat-anti-GFAP (Abcam, ab279291), Rabbit-anti-ARG1 (Thermo, PA529645), Mouse-anti-Aβ 6E10 (Biolegend, 803001), Rat-anti-MBP (Millipore, MAB386), Rabbit-anti-RTN3 (Millipore, ABN1723), Lamp1 (Biolegend, 121602).

Aβ plaques were visualized either by anti-Aβ 6E10 antibody or Methoxy X04 staining or by autofluorescence at 405 nm ^79^. For Methoxy X04 staining, sections were rinsed with ddH_2_O, and incubated with 34 μg/mL Methoxy-X04 (TOCRIS, 4920) diluted in 40% Ethanol for 15 min at room temperature, then rinsed twice with ddH2O and once with PBS, before subjecting to the blocking step.

For the APP/PS1:Cx43mgl-cKO and TAT-CX43@LNP experiment, mice were euthanized, and the brains were rapidly isolated, and separated into left and right halves. The left halves were used for acute brain slices experiments and the right halves were fixed by ice-cold PFA for immunostaining.

Images were captured using a VS200 slide scanner, FV3000 confocal microscope, or SpinSR spinning disk confocal microscope (Olympus). Superresolution images were captured using a SpinSR spinning disk confocal microscope.

### Microglia isolation and culture

Primary microglia were isolated and cultured as previously described ^80^ with minor modifications. Briefly, mouse pups were sacrificed on postnatal day 7. The brains were isolated, minced, and digested with 1 mg/mL papain for 1 hour at 37□. Tissues were then triturated with 10 mL pipette, and the resulting cell suspension was passed through a 70 μm cell strainer, and subjected to immunopanning with the rat-anti-mouse CD45 antibody (BD, 553076) and the secondary goat-anti-rat antibody (Jackson ImmunoResearch, 112-005-003) to isolate microglia. The immunopanned microglia were trypsinized and subjected to culture in microglia growth medium supplemented with TGF-β2/IL-34/Cholesterol and 1% fetal calf serum.

### Acute brain slice preparation and related experiments

The acute brain slices were prepared as previously described ^81^. Briefly, the mouse brains were rapidly isolated, stationed at the specimen holder, immersed in the ice-cold NMDG-based artificial cerebrospinal fluid (ACSF) with constant carbonation (95% O_2_, 5% CO_2_), and subjected to vibratome sectioning to obtain 280 μm slices. Slices were then recovered in NMDG-ACSF at 32 □, followed by sequential Na^+^ spike-in addition. The slices were then transferred to the carbonated ACSF at room temperature and recovered for at least 1 hour before experiments.

### Dye uptake

The dye uptake experiment was performed on acute brain slices as previously described ^24^. Acute brain slices were transferred onto 70 μm cell strainers put in the 6-well plate filled with the recording ACSF. Alternatively, CBX (200 µM), Gap26 (200 µM), and TAT-Cx43_266-283_ (50 μM) were added to the recording ACSF. After 15 min incubation, EtBr dye was added to the recording buffer at 5 µM, and incubated for 10 min. The slices were then rinsed 3 times with the recording ACSF (5 min each), fixed with 4% PFA for 1 hour, and subjected to immunostaining.

For the dye uptake experiment in primary microglial culture, cells were rinsed twice with Hank’s balanced salt solution (HBSS) (Ca^2+^/Mg^2+^ free), and preincubated in HBSS with or without TAT-Cx43_266-283_ for 15 min. As a negative control, CaCl_2_ was added to the HBSS at 1.8 mM. EtBr was then added to 5 μM, and further incubated with cells for 10 min. Cells were then fixed with 4% PFA for 10 min and subjected to microscopic analysis.

### Syncytial coupling - gap-FRAP

The gap-FRAP experiment was performed as previously described to analyze the connexin gap junction function ^82^. Briefly, acute brain slices were incubated with Sulforhodamine 101 (SR101) in the recording ACSF at 32 □ for 30 min. After rinsing with recording ACSF, slices were transferred to a recording chamber on the FV3000 confocal microscope. The baseline fluorescence of SR101 in the target cell was monitored for 45 s (1 frame per 5 s), followed by a 15-s photobleaching with 80% laser power in a 20 × 20 μm area centered at the target cell. The SR101 fluorescence was subsequently monitored for another 9 min to quantify the fluorescence recovery due to the SR101 dye diffusion from adjacent astrocytes through gap junctions.

### MitoSox staining

Acute brain slices were transferred onto 70 μm cell strainers put in the 6-well plate filled with the recording ACSF containing 5 μM MitoSox Red (Thermo, M36008), incubated for 20 min, then fixed with 4% PFA for 1 hour, and subjected to immunostaining.

### Supernatant collection and gliotransmitter detection

Acute brain slices were transferred to 1.5 mL microcentrifuge tubes containing 1 mL recording ACSF and incubated for 90 min. Supernatants were then collected for glutamate and D-serine measurement by commercial kits (Abcam, ab83389, ab241027), which were then normalized to the protein level of the slices measured by the BCA kit (Beyotime, P0012).

### TNF**α** and IL1**β** measurement

Acute brain slices were incubated with ice-cold lysis buffer (in mM, 50 Tris-HCl, 100 NaCl, 2 EDTA, 1% Triton-X100, supplemented with protease inhibitor cocktail), sonicated for 10 s, and centrifugated at 12000 g for 20 min. Supernatants were collected and stored at −80 □ before measurement. TNFα and IL1β level was determined by ELISA kits (Abclonal, RK00027 and RK00006), which were then normalized to the protein level measured by the BCA kit (Beyotime, P0012).

### RNA-sequencing

RNA-sequencing was performed on acute-isolated microglia from 9-month-old WT, APP/PS1, and APP/PS1:Cx43^mgl-cKO^ mice. Three biological replicates were included for each group. Mice were anesthetized and transcardially perfused with ice-cold HBSS. The brains were isolated with the meninges removed, followed by dissection of the cortex and hippocampus, which were then minced and digested by 1% papain. Microglia were isolated from the resulting cell suspension via immunopanning with the rat-anti-mouse CD45 antibody (BD, 553076) and the secondary goat-anti-rat antibody (Jackson ImmunoResearch, 112-005-003), and directly lysed on the panning plate by Trizol (Thermo, 15596018), and stored at −80□ before RNA extraction. RNA extraction and sequencing were commissioned to the Beijing Genomics Institute (BGI) using the DNBSEQ platform and analyzed using the HISAT-Bowtie2 pipeline. Differential expression analysis was performed on read counts using DESeq2. Pair-wise comparisons were performed between them. Differentially expressed genes were determined by a threshold of FC > 1.5, p < 0.05, and subjected to pathway enrichment analysis using DAVID (https://david.ncifcrf.gov/).

### TAT-Cx43@LNPs packaging and validation

TAT (YGRKKRRQRRR) and TAT-Cx43_266-283_ (YGRKKRRQRRR-AYFNGCSSPTAPLSPMSP) were synthesized by GL Biochem (Shanghai) Ltd. To package peptides into LNPs, 10 mg egg phosphatidylcholine (EPC) and 1 mg DOTAP were dissolved in 3 mL chloroform, followed by rotary evaporation at 40 □, 150 rpm for 1 hour to obtain a lipid thin layer. 30 mg Chitosan□Methacryloyl (CMA) was dissolved in 6 mL 0.9% NaCl and 0.01 M HCl to obtain a homogenous mixture, followed by the addition of 10 mg/mL photo initiator I2959 and 5 mg/mL peptides. The solution was then used to hydrate the thin layer of lipid for 1 hour. The resulting solution was filtered through 0.4 μm (3 times) and 0.2 μm (5 times) polycarbonate membranes (Whatman GE) to remove the multilamellar and unilamellar vesicles. The solution was then centrifuged at 300000 g for 1 hour to pellet the uncrosslinked nano lipid gel, which was then resuspended in 8 mL PBS (pH 7.4). The solution was then subjected to UV illumination (365 nm, 1-2 mW/cm^2^) for 30 min to induce polymerization, resulting in the DOTAP-NLG. The size and zeta potential of NLGs was then analyzed using Malvern Zetasizer 2000. To evaluate the release of peptides from NLGs, 1 mL sample was added to a dialysis bag immersed in 14 mL PBS, and shaken at 100 rpm, 37 □. The amount of peptide released into the PBS was then sampled and determined by spectrometry.

### TAT-Cx43@LNPs *in vivo* treatment and analysis

TAT-Cx43_266-283_ peptide was labeled with Cy5.5 before being packaged into LNPs. TAT-CX43@LNP or TAT-CX43 alone are i.p. injected into mice at 8 μL/g. Biofluorescence analysis was performed at different time points to analyze Cy5.5 distribution. At different time points post-injection, mice were anesthetized, followed by cardiac puncture and serum preparation, and perfused with ice-cold PBS. The brain, heart, liver, and kidney were then collected and lysed with RIPA buffer and sonicated for 10 s. Lysates were then centrifuged at 12000 g for 10 min, and the supernatants were collected. The Cy5.5 intensity was measured by a microplate reader, and the Cy5.5 intensity of the tissue lysate was normalized to the protein level measured by the BCA kit (Beyotime, P0012).

### ECG recording

Mice were anesthetized by i.p. injection of Tribromoethanol at 400 mg/kg. The ECG recording was then performed using the small animal ECG monitor (Xinrun, RM6240E). The data digitalization and analysis were performed with the RM6240E 2.5 software.

### Serum biochemistry analysis

Mice were anesthetized and ∼200 μL blood was collected via the retro-orbital sinus. The blood samples were then centrifuged at 12,000 g for 10 min, and the serum samples collected from the supernatant were stored at −80 □, until subjected to analysis using the Biostar V7 automated biochemistry analyzer (Seamaty, Ltd).

### Hippocampal LTP recording

Mice were anesthetized by isoflurane inhalation and transcardially perfused with iced-cold sucrose ACSF. The brains were isolated and stationed onto the specimen holder, immersed in carbonated ice-cold sucrose ACSF, and subjected to vibratome sectioning to obtain 300 μm thick brain slices. Brain slices were then transferred to ACSF and recovered at 32 □ for 1 hour, then at room temperature for 30 min before being subjected to LTP induction and recording. Slices were then transferred to the recording chamber with the ACSF flow rate of 1 mL/min. One monopolar electrode was positioned adjacent to the stratum radiatum of the CA1 region to stimulate the synaptic input and evoke field excitatory post-synaptic potential from the Schaffer collateral-commissural-CA1 synapse. For recording, another electrode was placed in the CA1 apical dendritic layer. The signals were amplified by a differential amplifier (Axoclamp 700B, Axon, USA) and were digitized using a Digidata 1550B (Axon, USA). An input-output curve was plotted before LTP induction. To set the test stimulus intensity, a fEPSP of 50% of its maximal amplitude was determined. Biphasic constant-current pulses were used for stimulation. LTP was induced using a theta-burst stimulation (TBS) protocol which consists of 50 bursts (consisting of 4 stimuli) at an inter-stimulus interval of 10 ms. The 50 bursts were applied for 30 s at 5 Hz (or at an inter-burst interval of 200 ms). The slopes of fEPSPs were monitored online. The baseline was recorded for 30 min. For baseline recording and testing at each time point, four 0.2 Hz biphasic constant-current pulses (0.1 ms/polarity) were used. fEPSPs were recorded every 5 min from 30 min before stimulation up to 240 min after stimulation across the CA1-CA3 Schaffer collaterals and normalized against t = 0. Data recording was performed using Clampex software, and data analysis was performed using Clampfit 10.8.

### Quantification

#### Quantification of cell numbers

The number of IBA1+ and GFAP+ cells was manually counted and normalized to the area of the region of interest. Only cells with visible cell bodies were included. To quantify the distribution of cells around the plaques, the distance between cells and the plaque center was measured for each plaque, and only cells within 100 μm of the plaque center were included; the cell-plaque distances were then sorted into 5-μm bins, which were then normalized to the plaque number quantified and then to the area within the 5-μm bin. To quantify the periplaque microglia number, the number of IBA1+ cells within 50 μm of a target plaque was manually counted, and normalized to the target plaque perimeter measured by Fiji.

#### Quantification of fluorescence intensity or positive area

Quantification of fluorescence intensity or positive area was performed using the Fiji software with or without macro scripts, where a threshold was set to distinguish the positive fluorescence signals, followed by the measurement of fluorescence intensity and positive area percentage within the region of interest. The same threshold was set for different experiment groups in the same batch. For images with high or uneven backgrounds, a subtract background function was performed with the rolling ball radius set at 50 pixels before quantification. To measure the fluorescence signal at different distances to the plaque, a serial of concentric circles with 5 μm or 1 μm escalating radius was drawn centering at the plaque center. The intensity or positive area was measured for each circle, and subtracted to the that of the smaller circle. To measure the number of RTN3 and Homer1 puncta at different distances from the plaque, a Sholl analysis was performed to quantify the intersection between signals and the concentric circle at different distances.

#### Signal masking

Masking of Cx43, Cx30, and Panx1 signals by IBA1 and GFAP was performed using the 3D counter function of the Fiji software. Masking of Methoxy X04 signal by IBA1 was performed by Imaris 9.0. The ratio of masked Methoxy X04 volume and the total Methoxy X04 volume was quantified, or plotted as XY plot to analyze the correlation. Microglia morphology was evaluated using the Sholl analysis plugin in Fiji software.

### Statistical analysis

Statistical significance between groups was determined with GraphPad Prism software. The unpaired t-test was used to determine the difference between two groups. One-way analysis of variance (ANOVA) was used to determine the difference among three and four groups. Two-way ANOVA was used to determine the difference between group variables in the curves. Pearson correlation test was used to determine the correlation between variables. Data distribution was assumed to be normal, but this was not formally tested. A probability of *p* < 0.05 was considered statistically significant. All significant statistical results are indicated within the figures with the following conventions: * *p* < 0.05, ** *p* < 0.01, *** *p* < 0.001, **** *p* < 0.0001. Error bars represent ±standard deviation. No statistical methods were used to predetermine sample sizes. The sample size per group was determined from previous publications using a similar methodology. Control mice and their littermate mutant mice were collected from multiple litters, and randomly selected for each experiment. Investigators were blinded to group allocation during data analysis. All experiments were performed at least three times, and the findings were replicated in individual mice and cell cultures in each experiment.

## Data availability

RNA-seq data have been deposited in the NCBI GEO database under the accession number GSE272781.

## Acknowledgement

Human tissue was provided by Human Brain Bank, Chinese Academy of Medical Sciences & Peking Union Medical College, Beijing, China, supported by the Neuroscience Center, Chinese Academy of Medical Sciences, and the Chinese Human Brain Banking Consortium. We thank Professor Arantxa Tabernero for providing expert advice on the TAT-Cx43_266-283_ peptide.

This work was supported by grants from the National Natural Science Foundation of China (NSFC 32170980 to C.Y.; 32271034 to J.N.; 32300791 to Y.S.), Chongqing Natural Science Fund for Distinguished Young Scholars (CSTB 2023NSCQ-JQX0030 to J.N.), National Key Research and Development Program of China (2021ZD0201703 to J.N.), Guangdong Basic and Applied Basic Research Foundation (2022B1515020012 to C.Y.; 2023A1515010817 to Y.S.; 2022A1515111165 to Q.W.), Science and Technology Planning Project of Guangdong Province (2023B1212060018 to C.Y.; 2021A1515110268 to Y.S.), and Shenzhen Fundamental Research Program (RCJC20231211090018040, RCYX20200714114644167, JCYJ20210324123212035, and ZDSYS20220606100801003 to C.Y.; JCYJ20210324121214039 and RCBS20210706092411028 to Y.S.; JCYJ20220530144816038 and RCBS20221008093118042 to Q.W.), Shenzhen Medical Research Fund (A2303014 to Y.S.).

## Supplementary Figure legends

**S Fig1.**
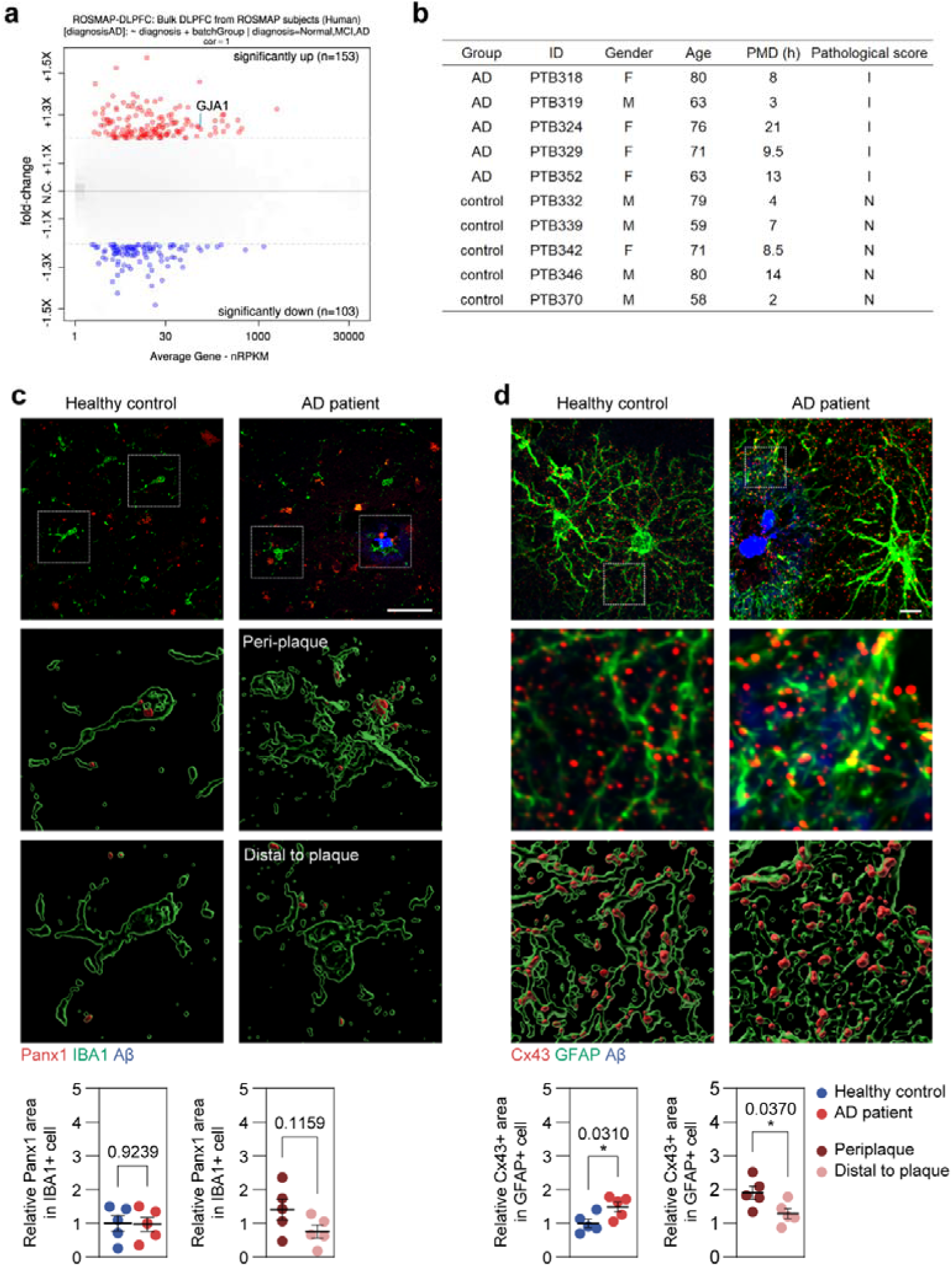
Cx43 expression in AD patients. (A) Volcano plot highlighting differential expressed genes in the ROSMAP-DLPFC-bulk RNA-seq dataset. Cx43/*GJA1* is among the upregulated genes in AD patients. (B) Information on human samples used in this project. (C) Immunostaining of Panx1, IBA1, and Aβ of human samples, and 3D reconstruction of Panx1 signals within IBA1+ domain. Scale bar: 50 μm. Below: quantification of Panx1+ area in microglia normalized to the controls. N = 5 subjects. (D) Immunostaining of Cx43, GFAP, and Aβ of human samples, and 3D reconstruction of Cx43 signals within GFAP+ astrocyte domain. Scale bar: 10 μm. Below: quantification of Cx43+ area in astrocyte normalized to the controls. N = 5 subjects. Dot plots show mean ± SEM, each data point, and *p*-value. Significance: * *p* < 0.05.

**S Fig2.**
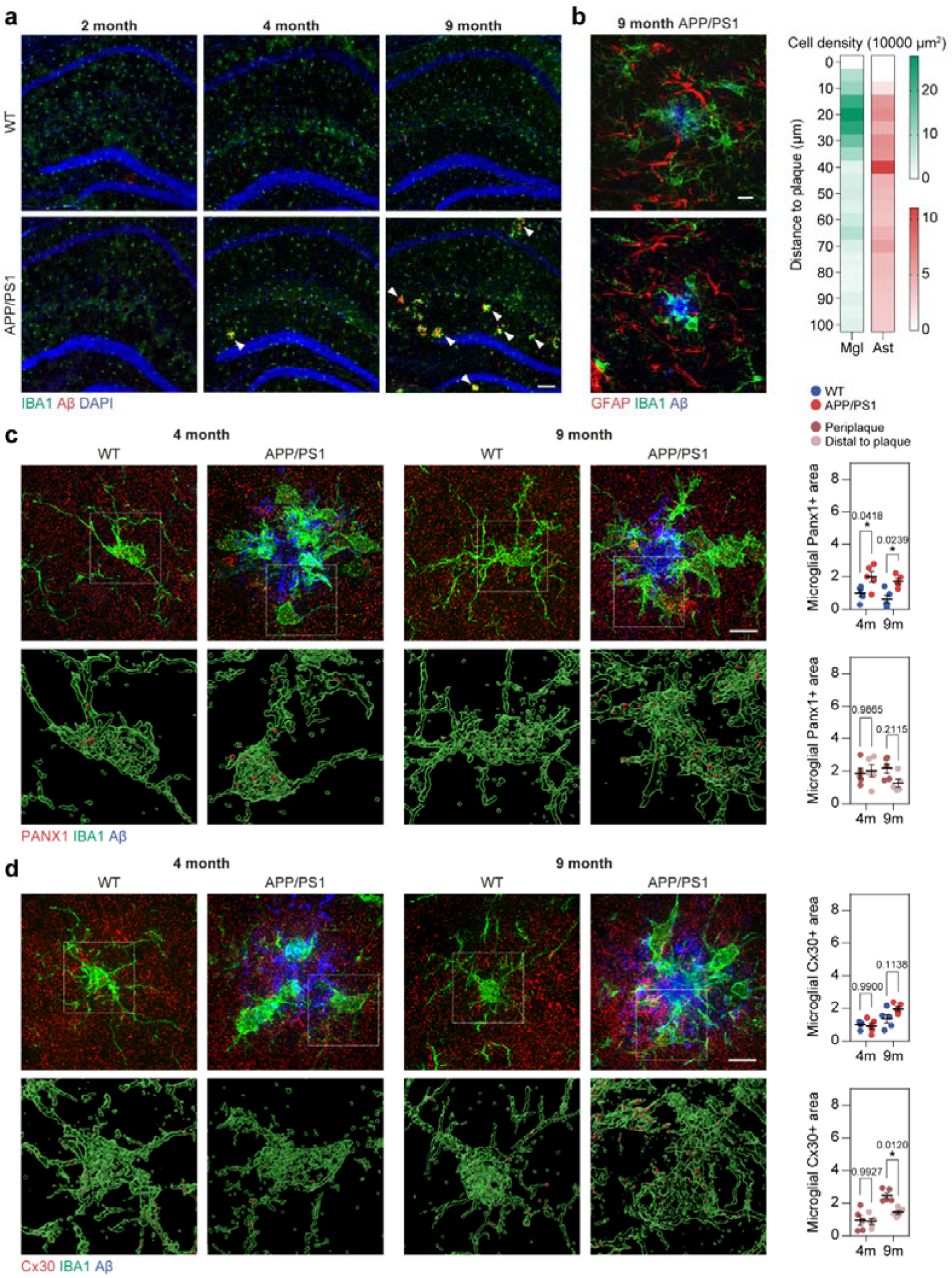
Microglial Cx43 expression and function in APP/PS1 mice. (A) Representative images of IBA1, Aβ, and DAPI staining in 2-, 4-, and 9-month-old WT and APP/PS1 mouse hippocampus. Arrowheads highlight Aβ plaque. Scale bar: 100 μm. (B) Representative images of IBA1, GFAP, and Aβ staining in 9-month-old APP/PS1 mouse hippocampus. Scale bar: 10 μm. Right, quantification of microglia and astrocyte distribution at different distances from Aβ plaques. Cells around a total of 15 plaques were calculated. (C) Representative super-resolution images of PANX1, IBA1, Aβ staining in 4-, 9-month-old WT and APP/PS1 mouse hippocampus, and 3D reconstruction of PANX1 signals within IBA1+ domain. Scale bar: 10 μm. (D) Representative super-resolution images of Cx30, IBA1, Aβ staining in 4-, 9-month-old WT and APP/PS1 mouse hippocampus, and 3D reconstruction of Cx30 signals within IBA1+ domain. Scale bar: 10 μm. Dot plots show mean ± SEM, each data point, and *p*-value. Significance: * *p* < 0.05.

**S Fig3.**
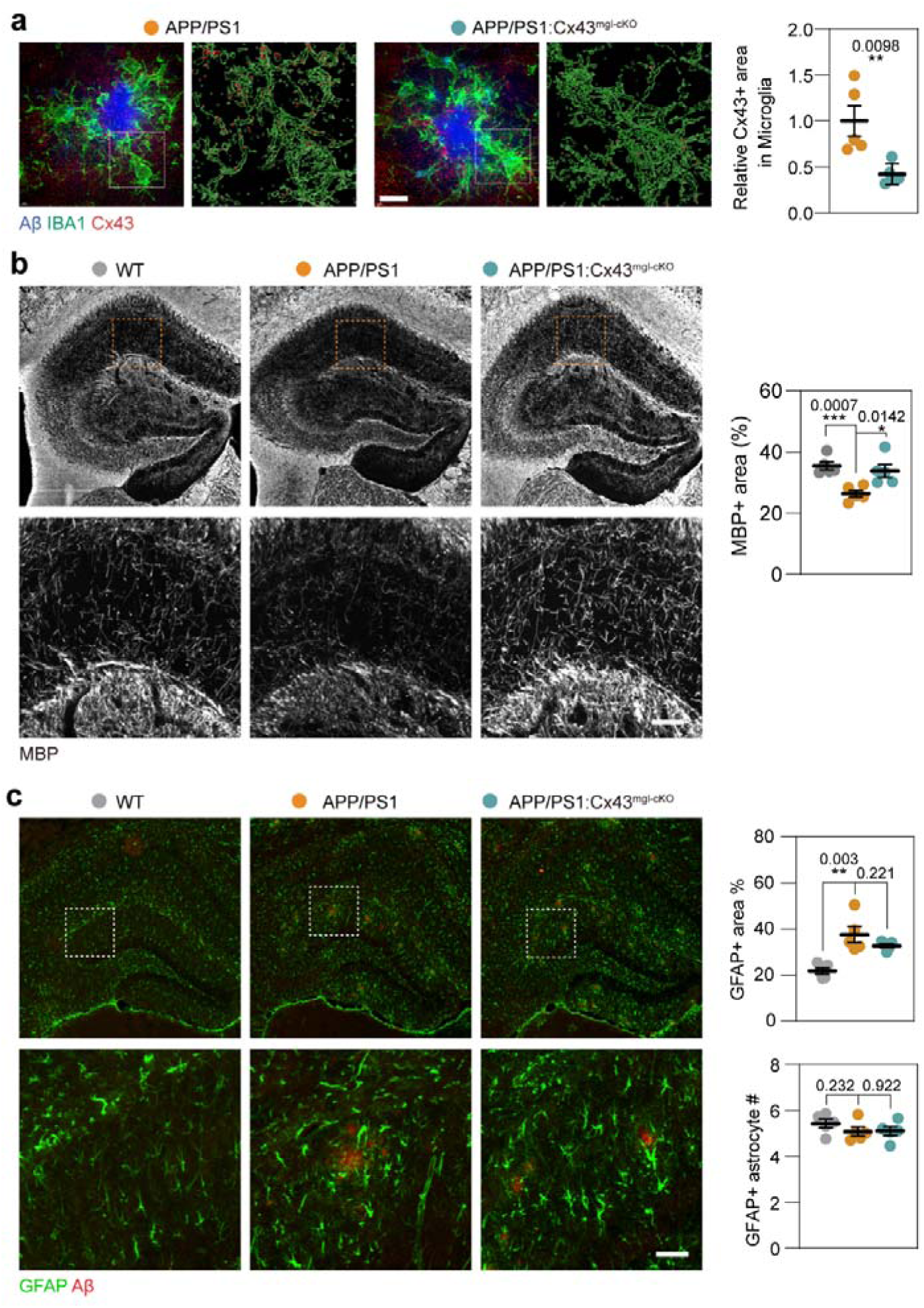
Histological analysis of Cx43^mgl-cKO^ in APP/PS1 mice. (A) Super-resolution image of IBA1, Cx43, and Aβ immunostaining of 9-month-old APP/PS1 and APP/PS1:Cx43^mgl-cKO^ mouse hippocampus, accompanied with 3D reconstruction of Cx43 signal within IBA1+ domain. Scale bar: 10 μm. Cx43+ signal area within microglia was quantified to verify the success of Cx43 knockout. N = 5 animals. (B) Representative image of myelin marker MBP staining. The MBP+ area was quantified. N = 5 animals. (C) Representative image of astrocytic marker GFAP staining. The GFAP+ area and GFAP+ cell number were quantified. N = 5 animals. Dot plots show mean ± SEM, each data point, and *p*-value. Significance: * *p* < 0.05, ** *p* < 0.01, *** *p* < 0.001.

**S Fig4.**
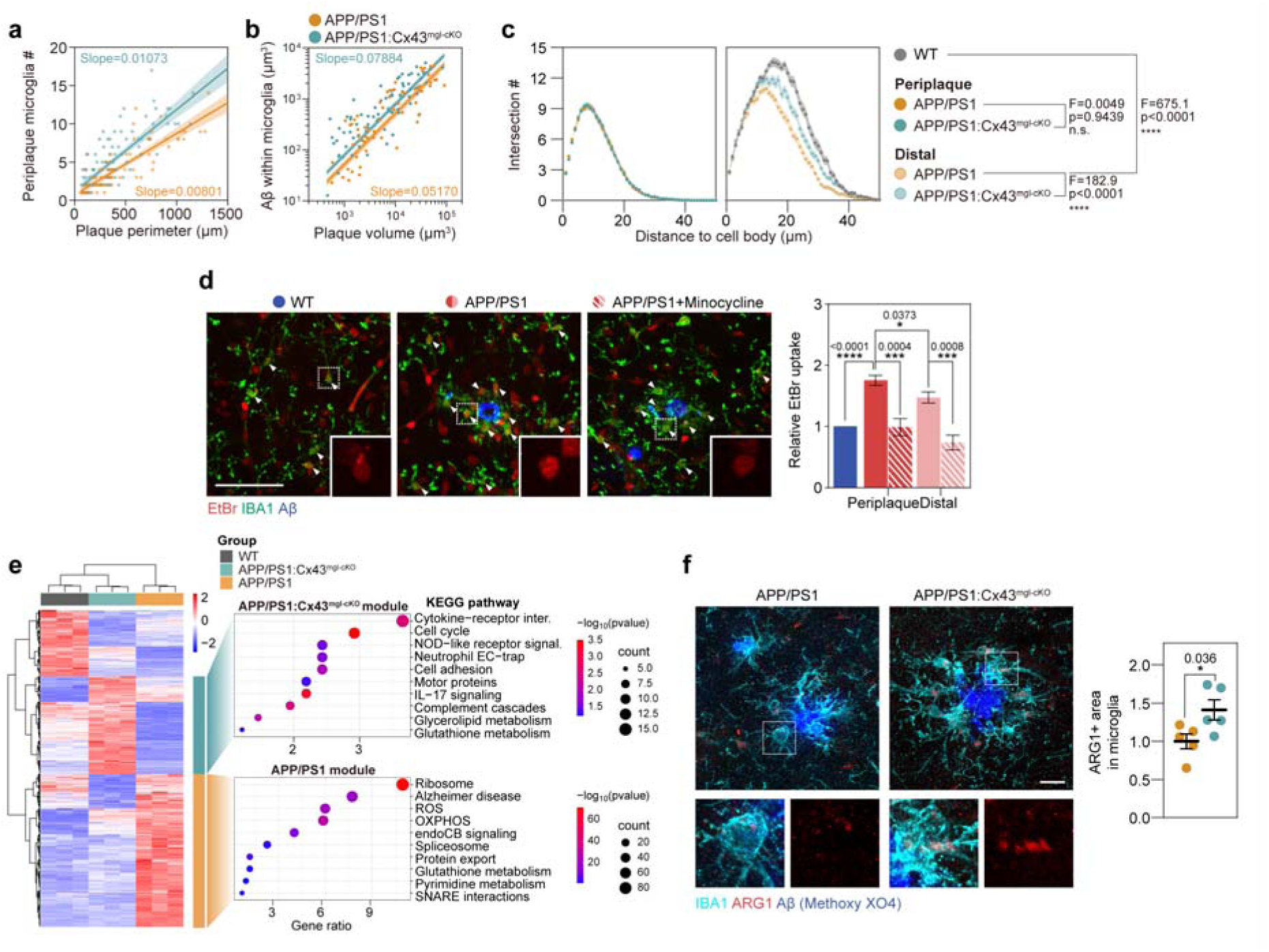
Cx43 ablation altered microglia reactive state in APP/PS1 mice. (A) The number of periplaque microglia of each plaque was plotted against the plaque perimeter, and the linear regression with 95% confidence interval was shown. N = 76 (APP/PS1), 106 (APP/PS1:Cx43^mgl-cKO^) cells. Related to Fig 4C. (B) The fraction of Aβ plaque within the microglia domain was plotted against plaque volume with the linear regression and 95% confidence interval shown. N = 63 (APP/PS1), 78 (APP/PS1:Cx43^mgl-cKO^) cells. (C) Left: Sholl analysis of periplaque microglia. Right: Sholl analysis of distal microglia and WT microglia. N = 83 (WT), 152 (APP/PS1-periplaque), 224 (APP/PS1:Cx43^mgl-cKO^-periplaque), 77 (APP/PS1-distal), 79 (APP/PS1:Cx43^mgl-cKO^-distal) cells. (D) Representative images of dye uptake experiment performed on 9-month-old APP/PS1 acute brain slices. Cx43-specific channel blocker GAP26 and pan gap junction/hemichannel blocker CBX were added to 9-month-old APP/PS1 acute brain slices. Scale bar: 10 μm. (E) Heatmap shows differentially expressed genes across WT, APP/PS1, and APP/PS1:Cx43^mgl-cKO^ microglia. Right: functional annotation analysis of APP/PS1 or APP/PS1:Cx43^mgl-cKO^ module, i.e., genes significantly enriched in APP/PS1 or APP/PS1:Cx43^mgl-cKO^ microglia. N = 3 animals. (F) Representative image of IBA1, ARG1, and Aβ staining. Arg1+ area in microglia was quantified and normalized to the mean of APP/PS1. N = 5 animals. Dot plots show mean ± SEM, each data point, and *p*-value. Significance: * *p* < 0.05, ** *p* < 0.01, *** *p* < 0.001, **** *p* < 0.0001.

**S Fig5.**
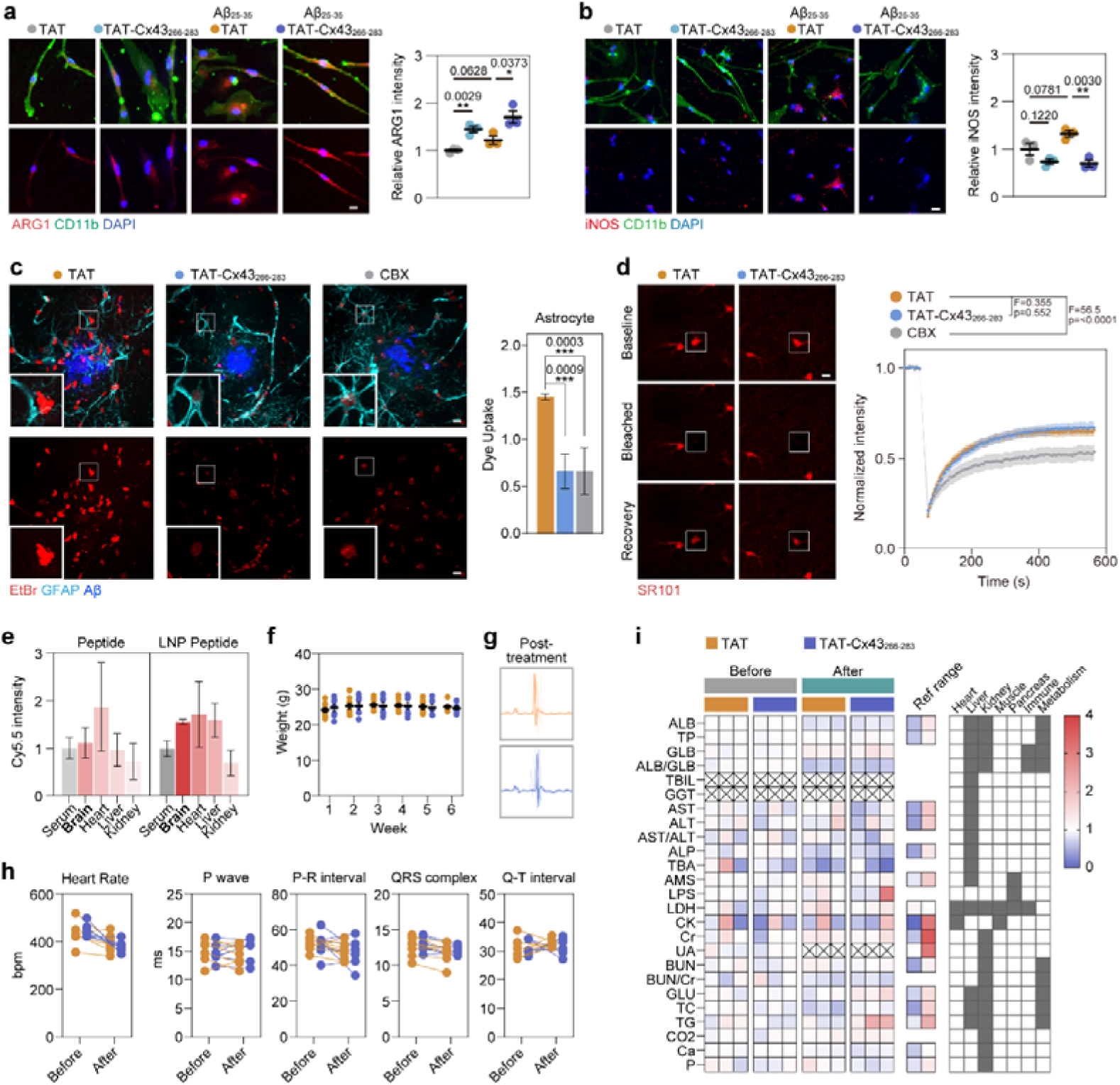
Evaluation of TAT-CX43@LNP. (A) Primary microglia were treated with 50 μM TAT or TAT-Cx43_266-283_ peptide and with or without 1 μM Aβ_25-35_ for 3 days before collection for immunostaining. Images show staining of ARG1, CD11b, and DAPI. Scale bar, 5 μm. ARG1 intensity was quantified and normalized to TAT-treated control. (B) Representative image of iNOS, CD11b, and DAPI staining of primary microglia. Scale bar, 5 μm. iNOS intensity was quantified and normalized to TAT-treated control. (C) Dye uptake experiment was performed on APP/PS1 acute brain slices and counterstained with GFAP. EtBr intensity of GFAP+ astrocytes was quantified. N = 4 experiments. (D) gap-FRAP experiment using SR101 dye on acute brain slices from APP/PS1 mice to examine the astrocytic gap junction function. Scale bar, 10 μm. (E) Tissue lysates were collected 1 day after TAT-Cx43_266-283_ peptide or DOTAP-NLG packaged TAT-Cx43_266-283_ peptide injection, and subjected to microplate reader analysis to analyze the tissue distribution of peptides. (F) Mice were weighed once a week during prolonged TAT@LNP or TAT-Cx43@LNP treatment. (G) Representative electrocardiogram of mice after 6-week treatment. (H) Quantification of Heart rate and electrocardiogram parameters before and after treatment. (I) Heatmap showing serum biochemistry parameters measured before and after TAT@LNP or TAT-Cx43@LNP treatments. Available reference ranges are listed, accompanied by the association between parameters and vital organs. ALB, albumin. TP, total protein. GLB, globulin. TBIL, total bilirubin. GGT, γ-glutamyl transpeptidase. AST, aspartate aminotransferase. ALT, alanine aminotransferase. ALP, alkaline phosphatase. TBA, total bile acids. AMS, serum amylase. LPS, lipase. LDH, lactate dehydrogenase. CK, creatine kinase. Cr, creatinine. UA, uric acid. BUN, blood urea nitrogen. GLU, glucose. TC, total cholesterol. TG, triglyceride. CO2, carbon dioxide. Ca, Calcium. P, phosphate. Dot plots show mean ± SEM, each data point, and *p*-value. Significance: * *p* < 0.05, ** *p* < 0.01, *** *p* < 0.001, **** *p* < 0.0001.

**S Fig6.**
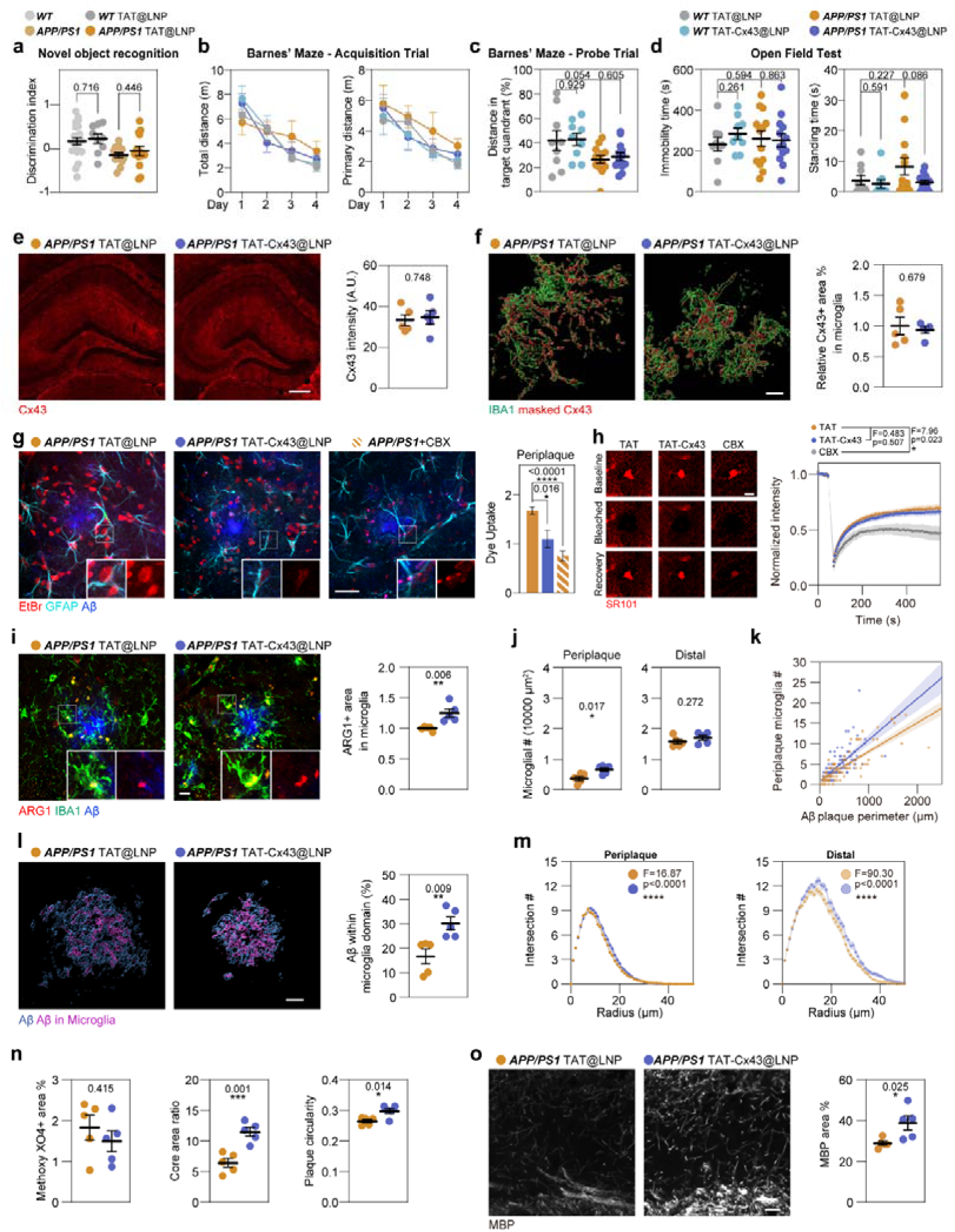
Effect of TAT-CX43@LNP on APP/PS1 mice. (A) Comparison of novel object recognition index between untreated animals and TAT@LNP treated animals. (B) Quantification of total distance, and primary distance to the target hole during Barnes’ maze acquisition trial. (C) Quantification of distance spent in the target quadrant during Barnes’ maze probe trial. (D) Quantification of immobility time, and standing time during the open field test. (E) Representative image of Cx43 staining of hippocampus and quantification of Cx43 intensity. Scale bar: 100 μm. N = 5 animals. (F) 3D rendering of IBA1 staining and Cx43 signal within IBA1+ domain. Scale bar: 10 μm. The microglial Cx43 signal was quantified, N = 5 animals. (G) Dye uptake experiment performed on acute brain slice followed by GFAP staining, Scale bar: 20 μm. The EtBr intensity of GFAP+ astrocyte was quantified. (H) Representative image of gap-FRAP experiments and quantification of SR101 dye intensity. Scale bar: 10 μm. (I) Representative image of IBA1, ARG1, and Aβ staining. Scale bar: 10 μm. Arg1+ area in microglia was quantified and normalized to the mean of APP/PS1. N = 5 animals. (J) Quantification of periplaque microglia and distal microglia normalized to the area. N = 5 animals. (K) the number of periplaque microglia was plotted against the plaque perimeter. The linear regression analysis was performed. (L) 3D rendering of Aβ plaque and Aβ plaque within IBA1+ microglial domain. Scale bar: 10 μm. The fraction of Aβ plaque within IBA1+ microglial domain was quantified. N = 5 animals. (M) Sholl analysis of periplaque and distal microglia. (N) Quantification of Methoxy X04+ Aβ plaque area, plaque core area ratio, and plaque circularity. N = 5 animals. (O) Representative image of MBP staining of hippocampal CA1 region. Scale bar: 10 μm. MBP+ area in the hippocampus was quantified. N = 5 animals. Dot plots show mean ± SEM, each data point, and *p*-value. Significance: * *p* < 0.05, ** *p* < 0.01, *** *p* < 0.001, **** *p* < 0.0001.

**S Fig7.**
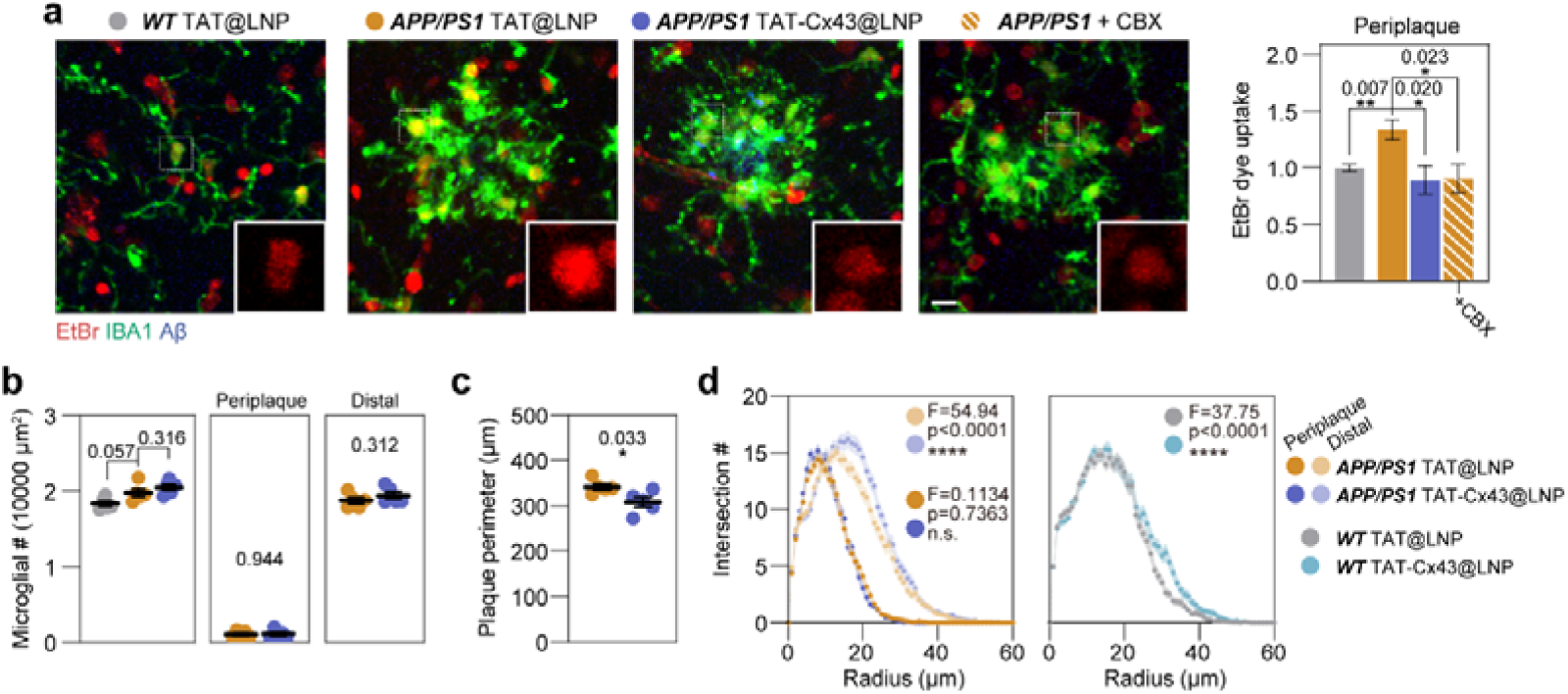
Early intervention with TAT-CX43@LNPs in APP/PS1 mice. (A) Dye uptake experiment was performed on APP/PS1 acute brain slices and counterstained with IBA1. Scale bar: 10 μm. EtBr intensity of IBA1+ microglia was quantified and normalized to WT. N = 5 experiments. (B) Quantification of the number of total microglia, periplaque microglia, and distal microglia normalized to the area. N = 5 animals. (C) Quantification of Plaque perimeters. N = 5 animals. (D) Sholl analysis of periplaque and distal microglia in APP/PS1 mice as well as WT microglia under TAT or TAT-CX43@LNP treatment. For periplaque microglia, N = 65 (TAT) and 72 (TAT-CX43) cells. For distal microglia, N = 50 (TAT) and 62 (TAT-CX43) cells. For WT, N = 42 (TAT) and 45 (TAT-CX43) cells. Dot plots show mean ± SEM, each data point, and *p*-value. Significance: * *p* < 0.05, ** *p* < 0.01, *** *p* < 0.001, **** *p* < 0.0001.

